# Lipid versatility of the C2C domain of Extended Synaptotagmin 3 safeguards plasma membrane binding

**DOI:** 10.64898/2026.10.13.758694

**Authors:** Saara Lautala, Theresa Jolie, Florian Wilfling, Dominik Oliver, Veronika Thallmair, Sebastian Thallmair

## Abstract

Membrane contact sites between the endoplasmic reticulum (ER) and the plasma membrane (PM) are critical hubs for lipid homeostasis and signal transduction. Extended Synaptotagmins (E-Syts) tether the ER to the PM and mediate lipid transport from the PM to the ER following Ca^2+^ signaling. Although E-Syts are essential for maintaining ER-PM contact sites during PI(4,5)P_2_ resynthesis, the molecular mechanisms governing their membrane association remain poorly understood. In particular, it is unclear how these contact sites are maintained following PI(4,5)P_2_ hydrolysis, when their principal membrane-binding ligand is transiently depleted.

Here, we characterize the membrane-binding mechanism of the C-terminal C2 domain of extended synaptotagmin 3 (E-Syt3) by combining coarse-grained molecular dynamics simulations with live-cell experiments. We identify distinct lipid binding sites with specific affinities for phosphoinositides as well as for phosphatidylinositol (PI) and phosphatidylserine (PS). While phosphoinositide interactions dominate membrane binding under resting conditions, PI and PS binding may provide an alternative membrane-binding mechanism that helps preserve ER-PM contacts during PI(4,5)P_2_ depletion. Our simulations further reveal that phosphoinositide binding is reinforced by the insertion of a tryptophan residue into the membrane hydrophobic core, providing a mechanistic basis for the high membrane affinity of the C2C domain. Together, our findings uncover the molecular determinants of E-Syt3 C2C-mediated membrane binding and provide mechanistic insight into the stability of E-Syt-mediated ER-PM contact sites during dynamic changes in membrane phosphoinositide composition.

## Introduction

Membrane contacts sites (MCSs) are sites of close membrane oppositions allowing for interorganelle communication, efficient intracellular signaling, and lipid trafficking.^1,2^ MCSs are formed by either protein tethers that bind simultaneously to both membranes or by a tethering pair of interacting proteins, each localizing to one of the opposed membranes.^3,4^ In this study, we investigate Extended Synaptotagmins (E-Syts), a protein family of endoplasmic reticulum (ER)-plasma membrane (PM) tethers that establishes ER-PM contact sites, recruits lipid transfer proteins, and also itself transports lipids between ER and PM.^5–8^

E-Syts are ER-resident and can dynamically bind the PM in trans through their multiple C2 domains.^5,9^ The E-Syt protein family consists of three homologues, E-Syt1, 2, and 3, all of which show the same basic structure depicted in Figure 1: ER binding is mediated by an N-terminal hydrophobic hairpin, which is followed by a lipid-transfer SMP domain and multiple C2 domains (three, termed C2A-C2C, in E-Syt2 and E-Syt3 and five, termed C2A-C2E, in E-Syt1).^5^ The C2A and C2B domains are arranged in tandem and are considered one membrane-binding module. In E-Syt1, C2A-C2B domains are evolutionally duplicated forming C2C-C2D. The most C-terminal C2 domains, C2C, in E-Syt2 and E-Syt3 are homologous to C2E in E-Syt1.^10^

**Figure 1:**
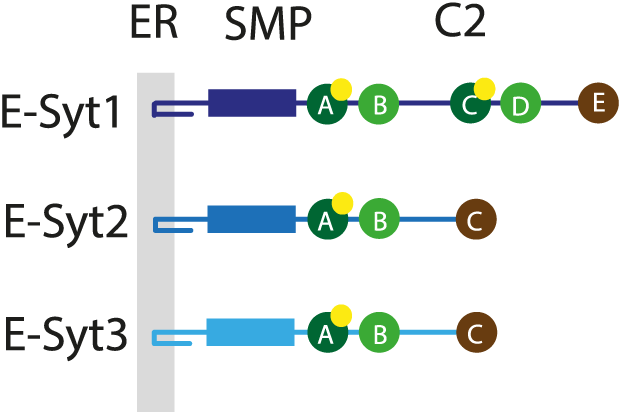
Domain structure of the Extended Synaptotagmin family members E-Syt1-3. E-Syts consist of an N-terminal ER-binding hairpin, a Synaptotagmin-like Mitochondrial-lipid-binding (SMP) domain, and multiple C2 domains. Color code of C2 domains indicates homology. Yellow circles indicate Ca^2+^ binding sites.

PM binding of all three E-Syt isoforms is phosphatidylinositol 4,5-bisphosphate (PI(4,5)P_2_) dependent and mediated by the terminal C2 domain (C2C in E-Syt2 and E-Syt3 and C2E in E-Syt1) and – in case of E-Syt1 – additionally by the central C2C domain.^5,9,11^ For E-Syt1, PI(4,5)P_2_ binding is Ca^2+^-dependent, resulting in an increase in tethering, as well as a tightening of the contact sites during conditions of high cytosolic Ca^2+^ concentration.^12^ Besides PI(4,5)P_2_ binding, the exact lipid binding properties of the individual C2 domains of E-Syt proteins as well as the molecular mechanism of membrane binding of the C2C domains are unknown. However, understanding the determinants of PM binding is of enormous functional significance, as this step not only initiates ER-PM contact formation but also determines the distance between the two membranes, which in turn regulates the recruitment of lipid transfer proteins as well as lipid transfer by the E-Syts themselves.^7,12^

Generally, E-Syts participate in the regulation of PM phosphoinositide homeostasis, ^6,7,12,13^ a family of signaling lipids of enormous functional significance. PM phosphoinositides are involved in numerous cellular processes including exo- and endocytosis, regulation of membrane proteins such as ion channels and transporters, and membrane dynamics. ^14^ They are generated from the common precursor PI, a lipid synthesized in the ER and transferred to the PM at ER-PM contact sites.^15,16^

Under resting conditions, E-Syt2 and E-Syt3 are PM bound, thereby establishing ER-PM contact sites of a width of approximately 19-22 nm.^17^ A rise in intracellular Ca^2+^ concentration, e.g. during PLCβ activation, additionally recruits E-Syt1 into the contact sites, which increases their number and cohesion but also decreases the membrane-membrane gap distance to approximately 15 nm. ^17^ However, it is not fully understood how the integrity of E-Syt tethered ER-PM contact sites is maintained and even tightened during PLC activation – and hence during PI(4,5)P_2_ depletion – as PM binding of all E-Syts is PI(4,5)P_2_ dependent. The decreased gap distance is a prerequisite for the recruitment of the PI transfer proteins, Nir2 and Nir3 into the contact sites.^12^ Besides providing PI to the PM, E-Syts are also involved in the clearance of PI(4,5)P_2_-derived second messengers like diacylglycerol (DAG) and phosphatidic acid (PA).^7,16^

Atomistic and coarse-grained (CG) molecular dynamics (MD) simulations are well-suited to study protein-lipid interactions.^18^ The balance between chemical accuracy, efficient sampling, and available lipid species renders the Martini force field a particularly suitable tool. ^19–21^ Using the Martini force field, lipid binding sites of transmembrane proteins were successfully identified, for instance cholesterol binding to G protein-coupled receptors^22^ and potassium channels,^23^ phosphoinositides to sodium channels, ^24^ and cardiolipin to complex III^25^ and bacterial proteins.^26^ A combined CG and atomistic MD pipeline enables the refinement of lipid binding sites in cryo-electron microscopy structures.^27^ Membrane binding was also identified for peripheral membrane proteins. For C2 domains, binding modes and orientation were investigated^28^ and their impact on the orientation of attached phophatase domains. ^29^ For Pleckstrin homology domains, well-known phosphoinositide binding domains, binding pockets were confirmed and concentration-dependent orientation^30^ as well as mutation effects were studied.^31^ In case of the Tubby domain, CG simulations identified PI(4,5)P_2_ binding sites and their cooperativity.^32,33^ Moreover, domains binding via cation-π interactions^34^ and hydrophobic patches ^35^ were resolved. Thus, CG MD simulations are a suitable tool to study the PM binding of E-Syts.

In earlier studies, E-Syt1 and E-Syt2 have received more attention than E-Syt3. Here, we embark on a journey to better understand the regulative role E-Syt3 holds at ER-PM contact sites. We start by taking a look at the interactions between the C2C domain of E-Syt3 and the plasma membrane. In doing so, we performed an in silico membrane binding assay at CG resolution using different membrane modes to characterize lipid preference, binding sites, and protein orientation on the membrane surface. Based on the in silico predictions, we performed live-cell mutation studies to characterize the membrane binding. Our joint computational and experimental study identifies previously unknown lipid binding sites of the C-terminal C2 domain of E-Syt3 which provide an unprecedented mechanistic understanding of its functional role for ER-PM contact sites.

## Results

### In silico identification of previously unknown protein-lipid interaction partners

The C2C domain of E-Syt3 is known to interact with the inner leaflet of the PM, specifically with PI(4,5)P_2_, however the exact binding site is not known. In addition, the role of other lipids in this interaction remains elusive. In order to explore these factors, we performed CG simulations with the Martini 3 force field^19^ containing a solvated C2C domain and different PM models (Figure 2a, for details see Materials and Methods). In total, six different simulation systems, each with a distinct membrane model, were set up to elucidate the lipid-specific interactions of the C2C domain. These membrane models represented the inner leaflet of a mammalian PM with varying levels of complexity, and their compositions were based on previous experimental and computational studies.^20,36–38^ Thus, the lipid species selected for our models were cholesterol (CHOL), 1-palmitoyl-2-oleoyl-*sn*-glycero-3-phosphocholine (POPC), 1-palmitoyl-2-oleoyl-*sn*-glycero-3-phosphoethanolamine (POPE), 1-palmitoyl-2-oleoyl-*sn*-glycero-3-phospho-L-serine (POPS), 1-palmitoyl-2-oleoyl-*sn*-glycero-3-phosphoinositol (POPI) and two different phosphoinositides, namely 1-palmitoyl-2-oleoyl-*sn*-glycero-3-phosphoinositol-(4,5)-bisphosphate (POPI(4,5)P_2_, hereafter PI(4,5)P_2_ or PIP_2_), and 1-palmitoyl-2-oleoyl-*sn*-glycero-3-phosphoinositol-4-phosphate (POPI(4)P, hereafter PI4P.) The six models were: 5MIX (28% CHOL, 25% POPC, 25% POPE, 18% POPS, 4% PIP_2_), 5MIX_doped_(27% CHOL, 24% POPC, 24% POPE, 17% POPS, 8% PIP_2_), 5MIX_PI4P_ (28% CHOL, 25% POPC, 25% POPE, 18% POPS, 4% PI4P), 5MIX_PI_ (28% CHOL, 25% POPC, 25% POPE, 18% POPS, 4% POPI), 4MIX (29% CHOL, 26% POPC, 26% POPE, 19% POPS) and POPC (100% POPC). Each of the six models and the lipid species are visually represented in Figure 2a. The lipid compositions were chosen to unravel the impact of different phosphoinositides within a simple model of the inner PM on the membrane binding of the E-Syt3 C2C domain.

**Figure 2:**
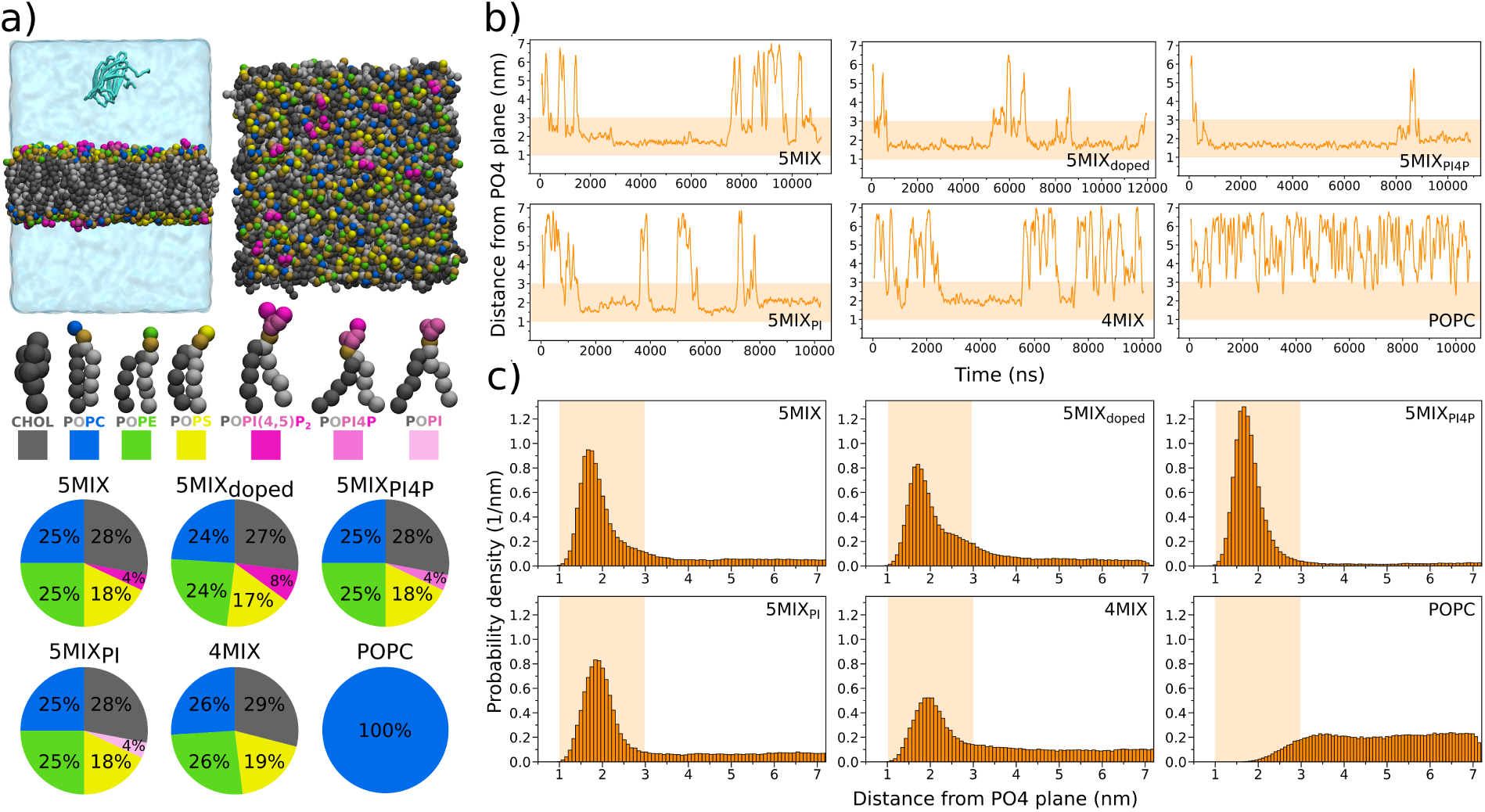
System setup and initial analysis. a) Graphical representation of the simulation setup and membrane components. Top panel shows a snapshot of the initial configuration of the 5MIX simulation box with the E-Syt3 C2C domain solvated in the water phase alongside an overhead view of the 5MIX membrane. Below these, each of the lipid species used in the simulations is represented as coarse-grained models, followed by the lipid compositions of the used membrane models. b) Plots demonstrating the dynamics of each systems via time-dependent series of the C2C domain’s distance to the membrane. The y-axis represents the z-distance between the center of geometry (COG) of the C2C domain and the phosphate bead plane (PO4 plane) of the nearest leaflet. Occupancy inside the highlighted region indicates contact between the C2C domain and the lipid bilayer. c) Normalized probability distributions of the C2C COG-PO4 plane distances over all repeats for each of the systems. Similarly to the time-dependent series, the highlighted region indicates the distance range for interaction with the bilayer. The observed hight of each of the peaks inside this region directly relates to the respective affinity of the C2C domain towards that membrane composition.

The simulations were first visually examined, revealing that in all systems except for pure POPC, the C2C domain interacted with the membrane in all repeats of the set. The interaction was quantified by calculating the time evolution of the z-distance between the center of geometry (COG) of the C2C domain and the closest phosphate bead plane (PO4 plane) of the membrane leaflets for all of the trajectories. Distances in the range of 1-3 nm (highlighted by orange rectangles) indicate a contact between the C2C domain and the lipid bilayer. Selected examples for each system are presented in Figure 2b, all other repeats are shown in Supplementary Figure S1 of the Supplementary Information (SI).

As expected from the simulation trajectories, the distance analysis for all mixtures excluding POPC, show substantial reversible interactions between the C2C domain and the respective membrane. There are multiple events of engaging and disengaging from the bilayer, with the engagements lasting up to several microseconds at a time. In case of the POPC membrane, the C2C domain has intermittent contact with the membrane surface, but no sustained interaction is observed. After time-dependent analysis, the distances observed from all trajectories of each system were combined into time-independent normalized distributions, presented as histograms in Figure 2c. Histograms for 5MIX, 5MIX_doped_, 5MIX_PI4P_, 5MIX_PI_ and 4MIX each clearly show a distinct maximum within 1-3 nm of the PO4 plane (highlighted by orange rectangles), indicating a highly populated membrane-bound state. For the POPC membrane, the distribution is flat in the water phase and tapers out towards the membrane indicating that there is no stabilizing interaction with the membrane, but that occasional contact of the C2C domain with the POPC bilayer results from diffusion as already observed by visual inspection.

Interestingly, the distributions for the membrane compositions are not identical; there is a deviation in both the binding affinity and the binding mode on the basis of the distance distributions. Firstly, the histograms show the probability densities of the interaction distance, and thus the hight of the histograms directly reflects the binding affinity between the C2C domain and the membrane in question. Out of the binding systems, 5MIX_PI4P_ has the highest maximum and thus strongest affinity, followed by 5MIX at second highest, then 5MIX_doped_ and 5MIX_PI_ at relatively comparable affinity, and lastly 4MIX with the lowest affinity. Secondly, the location of the maxima also varies depending on the system. 5MIX, 5MIX_doped_ and 5MIX_PI4P_ show a maximum position at around 1.75 nm, while for 5MIX_PI_ and 4MIX, this position is shifted towards 2 nm. Together, these differences imply that the C2C domain engages with the membrane in different manners depending on its lipid composition.

### PI(4,5)P_2_ and PI4P predominantly target C2C to the PM

We independently addressed phosphoinositide binding of the C2C domain experimentally in living CHO cells. Total Internal Reflection Fluorescence (TIRF) microscopy was used to image the association of the C2C domain to the PM while manipulating PM phosphoinositide levels. In detail, we co-expressed the fluorescently labeled C2C domain together with a system that allows stimulus-dependent recruitment of different phosphoinositide phosphatases to the PM. ^39,40^ Thus, a PM-anchored FRB domain was co-expressed with different FKBP-fused phosphatase constructs. FRB and FKBP hetero-dimerize in the presence of rapamycin resulting in the recruitment of the phosphatase constructs to the PM where they can exert their catalytic activity (Figure 3a). We either recruited the 4-phosphatase Sac1, which results in the selective depletion of PI4P, the 5-phosphatase Inp54, which depletes PI(4,5)P_2_, or pseudojanin, a synthetic 4- and 5-phosphatase that depletes both PI(4,5)P_2_ and PI4P (Figure 3b). As a negative control, we included a catalytically inactive mutant of pseudojanin (Ptase dead).

**Figure 3:**
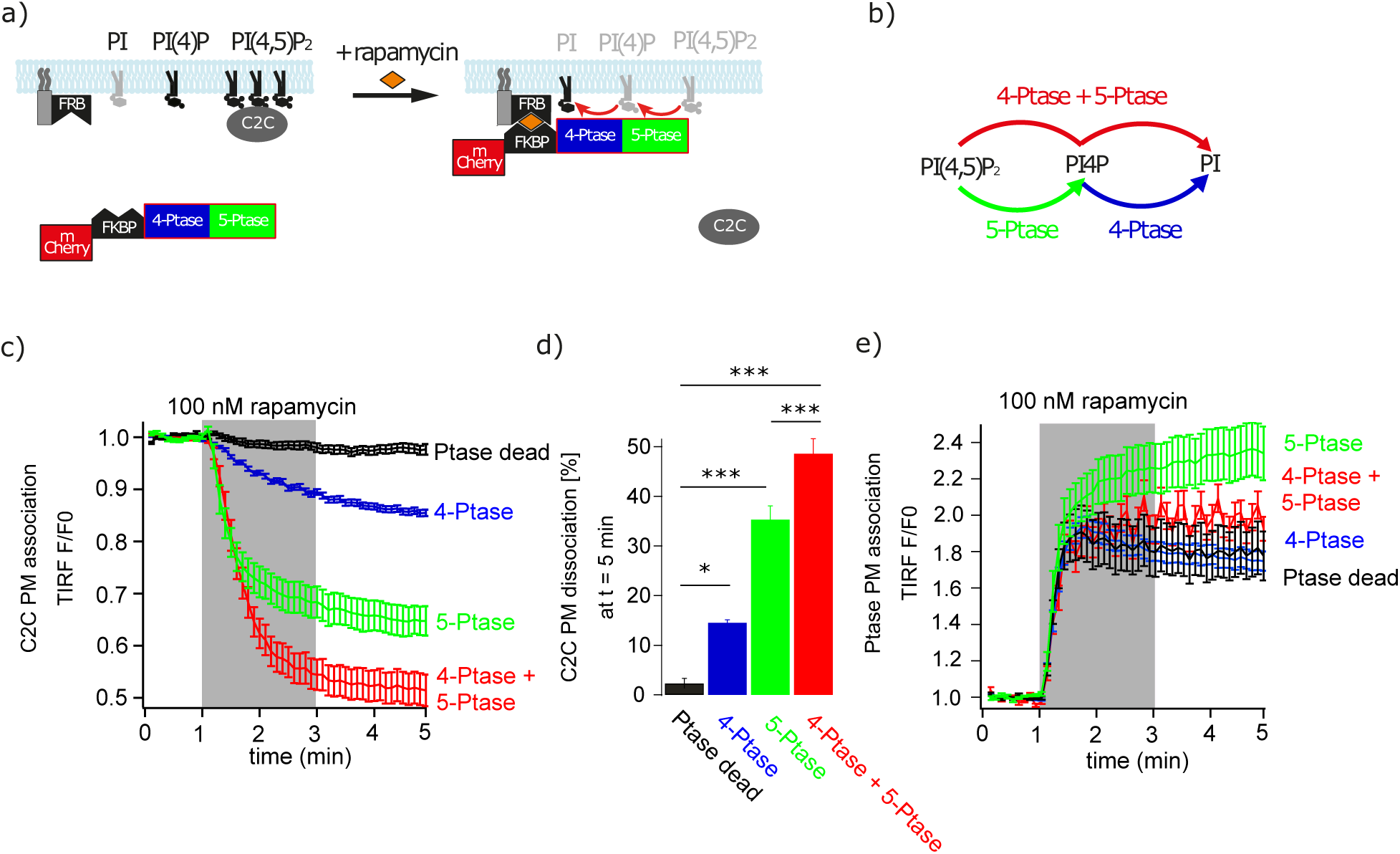
In the resting cell, the C2C domain binds to PI(4,5)P_2_ and PI4P. a) Scheme of the experimental set-up. Rapamycin-induced heterodimerization of FRB and FKBP induces PM recruitment of different FKBP-fused phosphoinositide phosphatases. Phosphatase recruitment to the PM as well as PM association of the C2C domain are tracked by TIRF microscopy. b) Catalytic reactions of used phosphatases. c) Normalized TIRF responses of the C2C domain upon recruitment of the various phosphatases shown in b. Depletion of PI(4,5)P_2_ but also of PI4P induced dissociation of the C2C domain from the PM. d) Statistical analysis of traces shown in c with *: p *<* 0.05, ***: p *<* 0.001. e) Relative TIRF fluorescences of recruited phosphatases derived from the same experiments as shown in c. Successful and comparable PM recruitment of the different phosphatases is confirmed.

All phosphatase constructs were successfully and comparably recruited to the PM (Figure 3e). The depletion of PI(4,5)P_2_ induced a strong dissociation of the C2C domain from the PM which is consistent with its known PI(4,5)P_2_ binding affinity^5^ also observed computationally (Figure 2c). Interestingly and further in line with our computational results, the depletion of PI4P also resulted in a significant detachment of the C2C domain from the PM, although to a smaller extent. The strongest dissociation was observed when both PI4P and PI(4,5)P_2_ were depleted simultaneously (Figure 3c,d).

These data support the computational observation that in addition to PI(4,5)P_2_, PI4P targets the C2C domain to the PM, a fact which may play physiologically an important role during conditions of PI(4,5)P_2_ depletion. However, the strong dissociation after Inp54 recruitment – which generates PI4P from PI(4,5)P_2_ – also emphasizes the preference of PI(4,5)P_2_ over PI4P binding at physiological conditions.

### In silico binding assay reveals lipid composition-dependent binding orientation of C2C domain

Upon visual inspection of the simulation trajectories, more than one face of the C2C domain was seen to associate prominently with the membrane. It became evident that the distance alone was not an accurate enough measure to elucidate the differences between the possible discrete binding modes, so we decided to additionally quantify the orientation of the C2C domain. To this end, a vector was defined for the C2C domain connecting the backbone beads of T782 and M830. To determine the orientation of the C2C domain with respeect to the membrane surface, the smallest angle between this vector and the membrane normal was calculated. A graphical illustration of the angle calculation setup is presented on the left panel of Figure 4.

**Figure 4:**
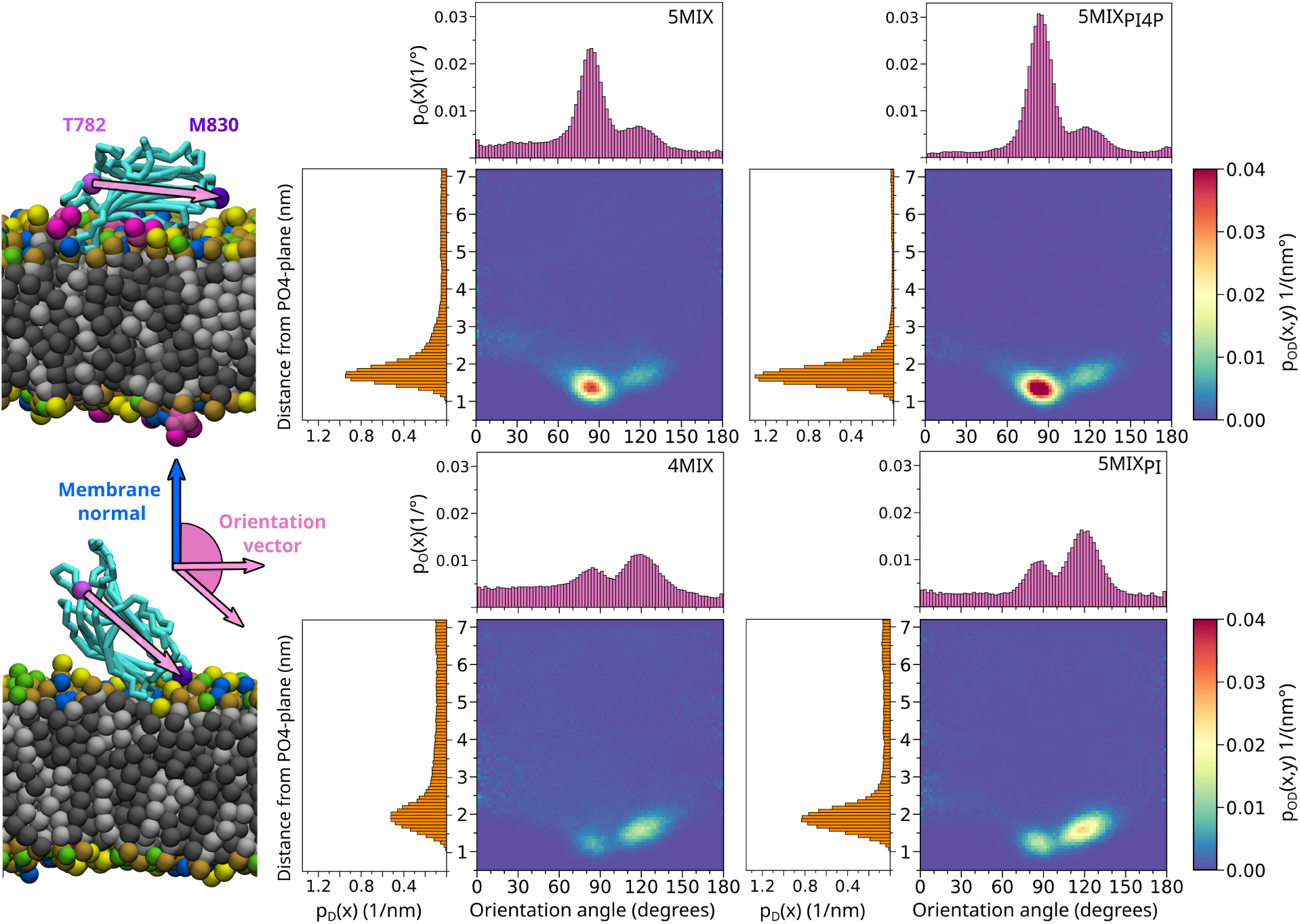
Orientation-distance population analysis of the C2C domain along with the visualization of the two dominant orientations, side binding (upper snapshot) and top binding (lower snapshot), on the membrane surface. First row shows the orientation-distance population profiles for 5MIX and 5MIX_PI4P_, while the second row shows these populations for the 4MIX and 5MIX_PI_ mixtures. Each 2D population is formed by combining the normalized distance and orientation angle probability densities, p_D_(x) and p_O_(x), which are presented on the left side and top of the 2D distributions, respectively.

Similar to the distance analysis, the time-dependent series were first evaluated individually for all repeats and then combined into time-independent normalized distributions. The orientation angle alone holds no information of membrane association. However, each measured value for angle is connected to a distance value for the same point in time. This correlation allowed us to combine the separate orientation and distance distributions into 2D orientation-distance distributions revealing the preferred binding populations for each lipid mixture. The results of this analysis for 5MIX, 5MIX_PI4P_, 5MIX_PI_, and 4MIX are presented in Figure 4, the distributions for 5MIX_doped_ and POPC are depicted in Supplementary Figure S2 in the SI.

The results clearly show two distinct populations, one around distance 1.75 nm and orientation angle *∼*85*^◦^*(hereafter ‘side-binding’ orientation), and one around 2 nm and *∼*120*^◦^*(hereafter ‘top-binding’ orientation). Both populations are present in all membrane compositions, but it is evident that the side-binding orientation dominates in the 5MIX, 5MIX_doped_ (Supplementary Figure S2), and 5MIX_PI4P_ membranes, while the top-binding orientation dominates in the 4MIX and 5MIX_PI_ membranes. For the POPC membrane, the 2D distribution does not show any preferred orientation which is consistent with the lack of membrane binding already observed in the distance analysis above (Supplementary Figure S2).

From this analysis it is clear that there are two distinct binding orientations for the C2C domain, and the predominant population is directly related to the lipid composition of the relevant membrane. The analysis indicates that mixtures with phosphoinositide lipids (PIPs), prefer the side-binding orientation, while systems with less negatively charged lipids, such as PS or PI, prefer the top-binding orientation.

### Different lipid species exhibit individual binding sites

Based on the lipid-dependent orientation of the C2C domain on the membrane surface, we hypothesized that different lipid species might have individual binding sites which ultimately result in the observed differences in orientation. In order to identify the key residues which mediate the interactions with the membrane for each of the binding modes, we analyzed the number of lipids which are in contact with the individual residues of the C2C domain. Figure 5 shows the averaged contact analysis for 5MIX, and 4MIX. These two mixtures representatively illustrate this difference in lipid binding. The data for the other lipid mixtures are depicted in Supplementary Figure S3 in the SI.

**Figure 5:**
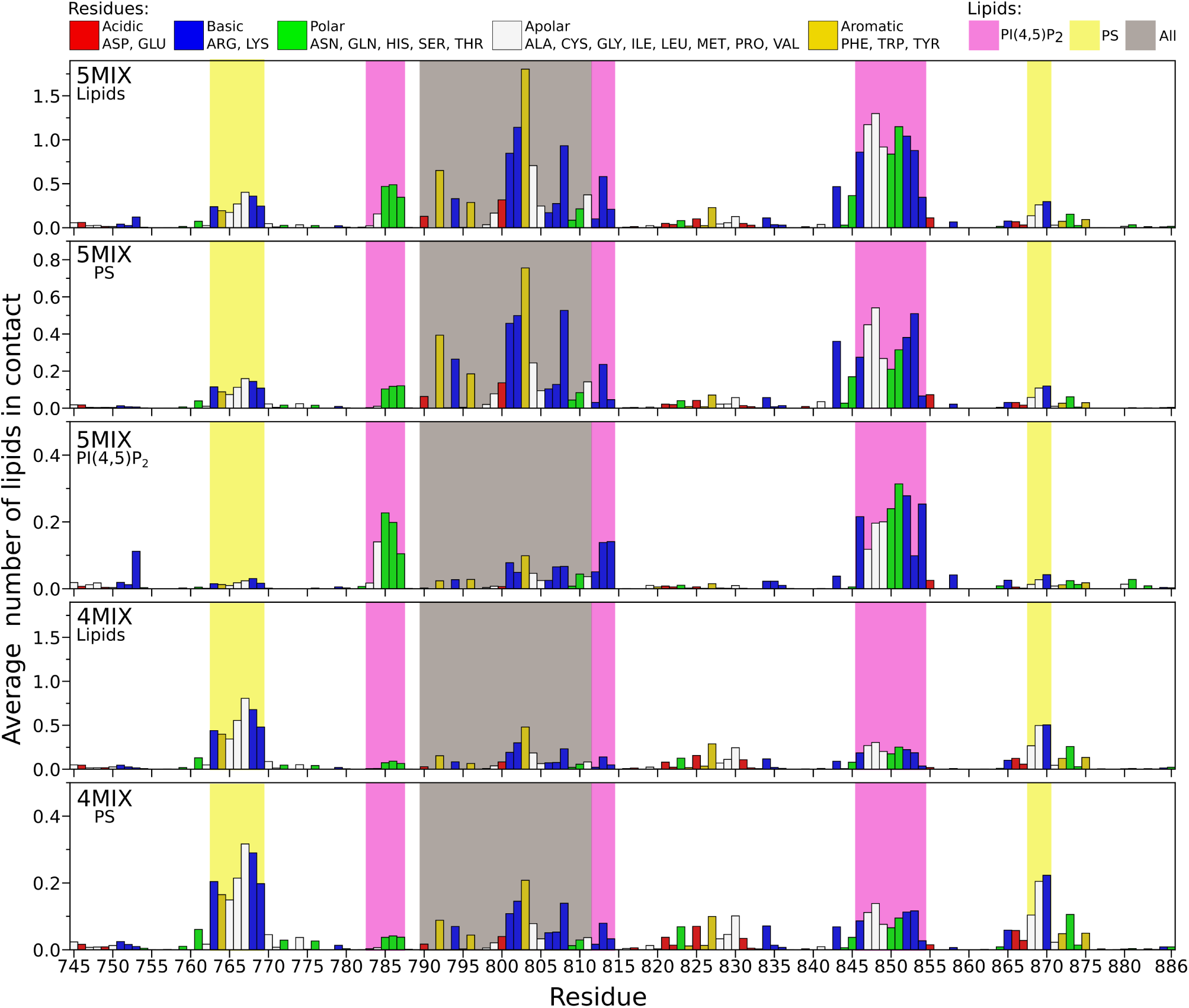
Average contacts with the bilayer per residue plotted as a function of the C2C-domain sequence for 5MIX and 4MIX systems. Amino acids are colored by their chemical properties, with red representing acidic (negatively charged at physiological pH), blue for basic (positively charged at physiological pH), green for polar, white for apolar, and mustard yellow for aromatic. The first panel shows average contacts for all lipids in 5MIX, while the second and third show contacts for specifically PS and PI(4,5)P_2_, respectively. The last two panels show average contacts for all lipids and PS, respectively, in the 4MIX system. Regions highlighted with magenta indicate PI(4,5)P_2_ and yellow for PS interaction enrichment. Please note that for visibility, the y-axis scaling is not uniform across all the panels.

When comparing the plots for 5MIX and 4MIX, there are multiple regions enriched in lipid interactions for both lipid mixtures. Firstly the region between residues 763-770 (highlighted yellow) is clearly enriched in lipid interaction in both mixtures. This region is specific to PS, as PS interaction in both mixtures makes up most of the total interactions, and in 5MIX there is virtually no interaction with PI(4,5)P_2_ in this region. Similarly, a region at around residues 868-870 (highlighted yellow) in both mixtures shows preferred interaction with PS. With respect to the total lipid interactions (first and fourth panel), both regions are more prominent in 4MIX than in 5MIX, which indicates that these regions are associated with the top binding mode.

Simultaneously, the region between residues 845-855 (highlighted magenta) also shows increased interaction in both mixtures, but it interacts preferentially with PI(4,5)P_2_ rather than with PS. Interactions of all lipids with this region are clearly enriched in 5MIX compared to 4MIX. Closer inspection shows that even though both PI(4,5)P_2_ and PS interact with this region in 5MIX, the interaction of PS in 4MIX is considerably weaker. There are also smaller patches of interacting regions around residues 783-787 and 812-814 (both highlighted magenta) showing up in 5MIX but not in 4MIX, indicating a preference for PI(4,5)P_2_. We hypothesize that these regions drive PM association in the side binding mode in 5MIX.

Interestingly, in 5MIX but not in 4MIX, there is also a rather large region of high interaction around residues 790-811 (highlighted taupe), which is rather unspecific with respect to the lipid species. Neither PI(4,5)P_2_ or PS show significant enrichment in this area, and furthermore, the residue that interacts the most in this region is W803, a hydrophobic aromatic residue. This indicates that this region can interact with the hydrophobic core of the membrane. The stronger interaction of this region in 5MIX than in 4MIX indicates that it likely relates to the side-binding orientation, and not the top-binding orientation.

The comparison between 5MIX and 4MIX helped us to recognize clear interaction sites specific to each of the binding orientations in the protein sequence. Next, we projected the results of the contact analysis to the atomistic protein structure to see the spatial proximity of the distinguished areas within the folded protein, and to potentially relate them to the results of the distance-orientation analysis. Figure 6 depicts the C2C domain colored according to the average lipid contacts for 5MIX, 4MIX, 5MIX_PI4P_, and 5MIX_PI_. The first row shows the average contacts for all lipids. Remarkably, the contact patterns can be divided into two groups, namely the mixtures linked to side-binding orientation, *i.e.* 5MIX and 5MIX_PI4P_, and the ones linked to the top-binding orientation, 5MIX_PI_ and 4MIX. While for 5MIX and 5MIX_PI4P_ the majority of the contacts reside at the two loops at the bottom of the structure (residues 790-811 and 846-854), in 4MIX and 5MIX_PI_ the interactions are concentrated on the right side of the structure at the end of the principal axis of the protein (residues 763-770 and 868-870). When looking at the lipid species-specific interactions presented in the second row of Figure 6, the same trend persists. Moreover, for the side-binding mixtures, two additional loops located further from the membrane (residues 784-787 and 812-814) participate in the PIP interaction on the left side of the structure. While the lower PIP interaction loop is intermittently interacting also in mixtures without PIP, contacts with the additional loops are absent (see both Figures 5 and 6). It is likely that these residues are only accessible for PIPs due to their larger headgroups, even in the side binding orientation. This suggests, that the specificity of the PIP and PS/PI binding sites likely is not solely caused by the difference in the charge of the interacting lipid species, but also their respective geometries and sizes.

**Figure 6:**
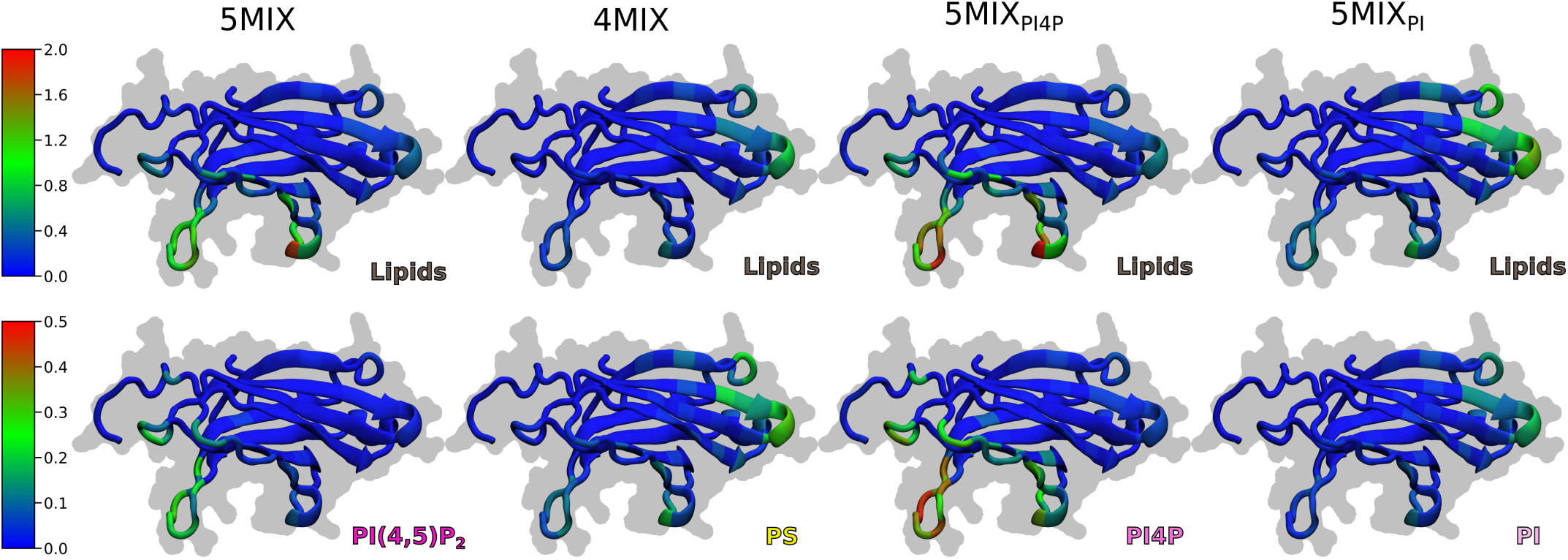
Average contacts with the bilayer per residue mapped on the folded structure of the C2C domain. The first row shows average contacts with all lipids in the respective system, while the bottom row shows contacts with a selected specific lipid species in that system. The colorbar for each row is visible on the left, with red always indicating high amount of contacts and blue low amount of contacts.

Taken together, PIP lipids drive the interaction with 5MIX and 5MIX_PI4P_ in the side-binding orientation, while PS and PI, respectively, drive the interaction with 4MIX and 5MIX_PI_ in the top-binding orientation.

### The PIP binding site is the preferred binding site under resting conditions

To test whether the PIP and PS/PI binding sites affect the resting PM localization of the C2C domain in living cells, we over-expressed various GFP-tagged C2C mutants together with the PM marker L10-RFP in CHO cells. We performed life-cell confocal imaging and analyzed the relative PM localization (see Section Confocal microscopy) of the mutants. The mutants were generated on the basis of the lipid interaction profiles identified in the MD simulations (Figure 6). We generated single mutants and double mutants of the PS/PI and of the PIP binding sites, respectively, as well as two mutants that contained mutations in both binding sites. The mutants of the PS/PI binding site were R763A and K870A single mutants, as well as R763A K870A and R768A R769A double mutants. At the PIP binding site, we generated single mutants R813A, K814A, R846A, R852A, R853A, R854A and double mutants R813A K814A and R852A K854A.

The wild-type C2C domain, as well as the PS/PI binding mutants showed robust co-localization with the PM marker, which can be seen in example images (Figure 7c and Supplementary Figure S5 in the SI). For quantification, relative PM localization was determined from line profiles drawn across an entire cell. Relative PM localization was calculated as the ratio of the fluorescence at the PM peaks (defined by the PM marker peaks) and the cytosolic fluorescence (Figure 7c). Relative PM localization of the PS/PI binding site mutants reached values of approximately 2 and was not significantly different from that of the wild-type domain. In contrast, the single mutants R813A, K814A, and R854A as well as the double mutants of the PIP binding site showed significantly reduced PM localizations (Figure 7a,c).

**Figure 7:**
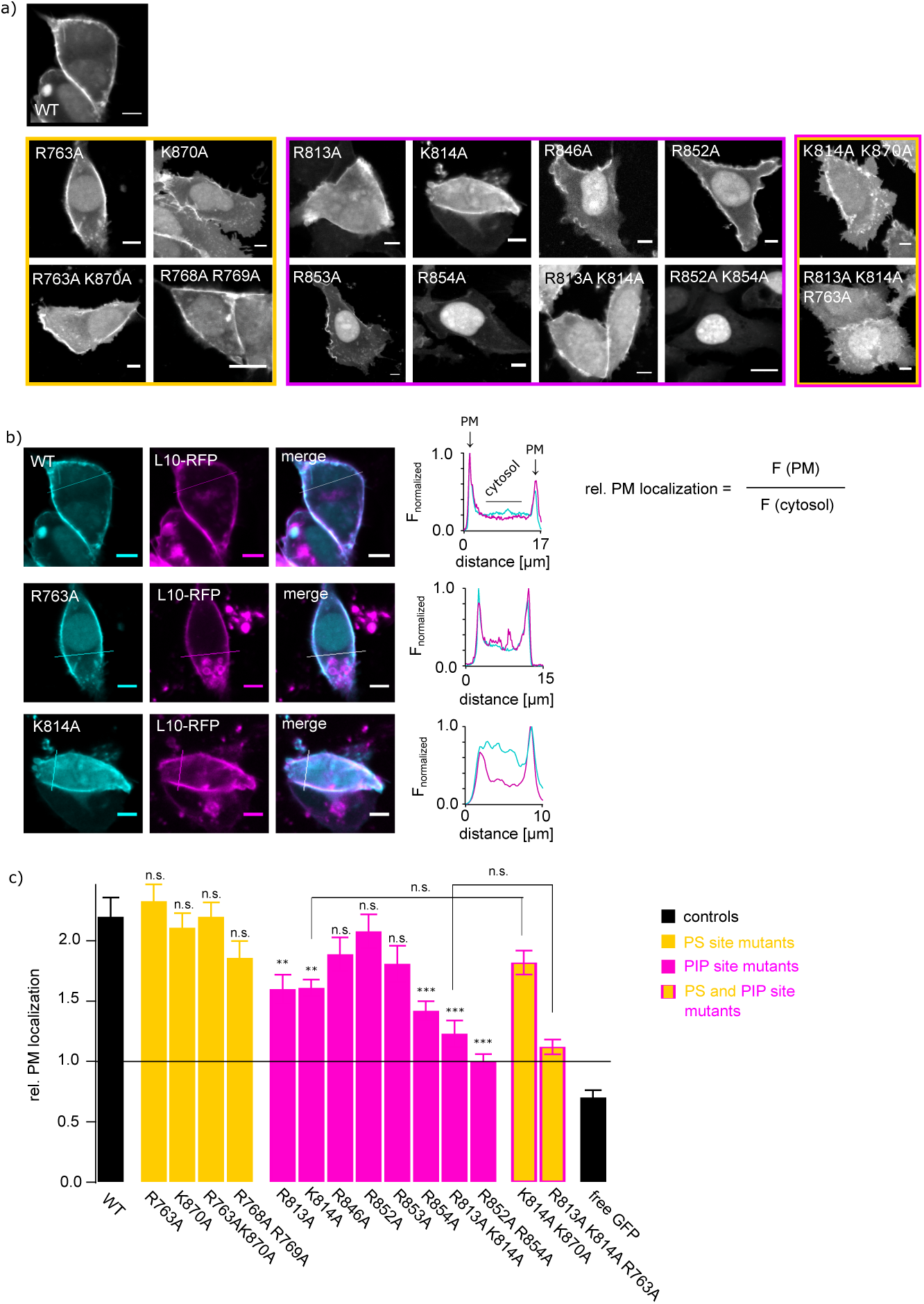
Under resting conditions, the PIP binding site determines PM localization of the C2C domain. a) Confocal images showing the localization of C2C wild-type domain and various mutants of the PS (yellow) and PIP (purple) binding site, respectively. CHO cells were transiently transfected with GFP-tagged C2C variants and RFP-tagged PM marker L10. Representative confocal images of GFP fluorescence are shown. Scale bar = 5 µm. b) PM localization analysis of GFP-tagged C2C domains (wild-type and two example mutants). Images derive from experiments as in a. GFP and RFP fluorescence is shown in cyan and magenta, respectively. Right: Normalized fluorescence intensity line profiles from highlighted lines. Relative PM localization was calculated from normalized line profiles. Note: Reduced PM localization of K814A mutant can be seen in an increased cytosolic fluorescence intensity in the normalized line profile. c) Relative PM localizations of all investigated mutants and controls. Values derived from line profile analysis as shown in b. Dunnetts testing revealed significant reduced PM localization of several mutants of the PIP binding site compared to the wild-type PM localization (**: p *<* 0.01; ***: p *<* 0.001).

We might have overlooked an effect of the top binding site mediated by the PS/PI binding site, as the stronger PIP binding site may have taken over in the top binding mutants. Thus, we also generated two mutants K814A K870A and R813A K814A R763A with mutations in the PIP and the PS/PI binding sites. The PM localization of these two mutants was not significantly different from the localization of the respective mutants of the PIP binding site only (see Figure 7c and Supplementary Figure S5 in the SI), indicating that the PS/PI binding site does not significantly contribute to the PM localization of the C2C domain, at least under resting conditions.

**Occupancy of binding sites determines orientation on the membrane.**

Both the side-binding and the top-binding orientations are present in the distance-orientation analysis for all mixtures which show significant membrane binding. However, based on the contact analysis, the interacting sequences lie spatially far apart in the 3D protein structure. This implies that their simultaneous engagement is unlikely on a flat membrane, and we hypothesized that the systems alternate between the two orientations during the simulations. To verify this, we defined simplified discrete binding regions on the residue level based on the identified binding sites and compared the time evolution of their interactions with the lipids during our simulations. The defined binding regions focused on the most strongly interacting residues for the PIP and PS/PI interacting regions, and W803 was selected as representative for the hydrophobic region to explore the contribution of the hydrophobic region to the observed binding poses. The PIP binding site contains two distinct areas in the structure-mapped contact analysis (see Figure 6). This lead to the question, whether these loops act collectively or separately during the binding. To address this question, we divided the PIP binding site in two separate sub-regions for this analysis, PIP I being closer to the membrane and PIP II being located slightly above the membrane in the side-binding orientation. The residues of the analyzed binding regions are given in Table 1.

**Table 1:** Residues of the C2C domain which are part of the different binding regions analyzed in. **Figure 8**. Residues in bold font were tested experimentally in different combinations by means of mutation experiments.

| Binding site | Selected residues |
| --- | --- |
| PIP I | R753, **R846**, S850, H851, **R852**, **R853**, **K854** |
| PIP II | T785, S786, S787, **R813**, **K814** |
| PS/PI | **R763**, C766, L767, **R768**, **R769**, I869, **K870** |
| HydPho | W803 |

Figure 8 depicts the time evolution of the number of lipids within a distance of 0.6 nm of the respective binding regions for the 5MIX, 5MIX_PI4P_, 5MIX_PI_, and 4MIX systems together with the time evolution of the protein orientation angle for one repeat of each simulation series.

**Figure 8:**
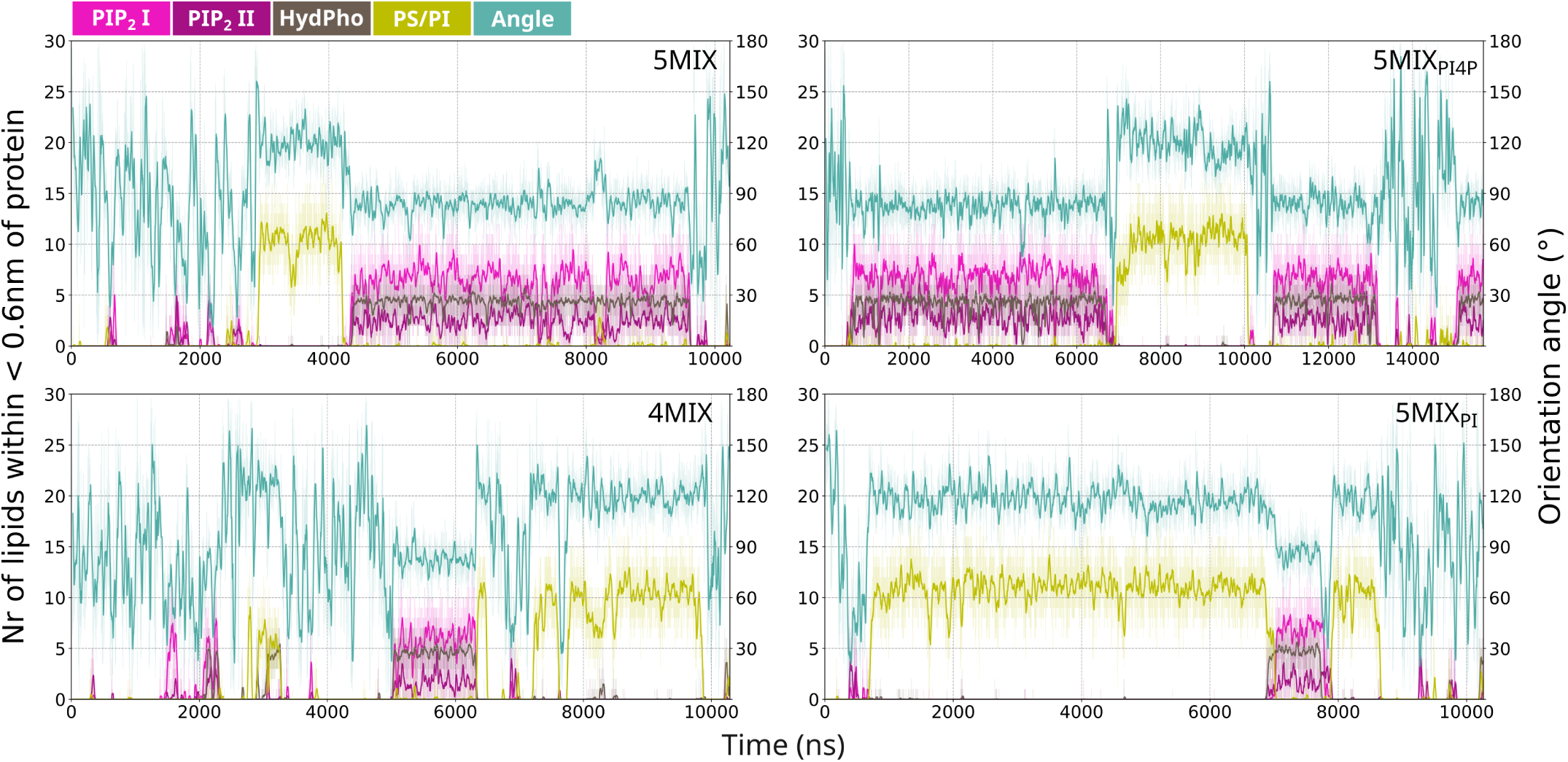
Temporal evolution of the lipid interactions of the binding sites defined in Table 1 and protein orientation angle. The transparent lines represent the raw data with 250 ps interval, while the bold lines represent the running average of the data with a 25 ns sampling window.

All replicas are depicted in Supplementary Figure S4 in the SI. It can be easily recognized that the C2C domain alternates between two binding modes during the simulations – either being in contact with the PS/PI binding region resulting in the top-binding orientation or being in contact with the PIP I, PIP II, and the hydrophobic binding region resulting in the side-binding orientation. As expected, there is no significant temporal overlap of the contacts between the PS/PI and PIP regions, clearly showing that the two binding regions do not interact simultaneously. Interestingly, there is also no significant temporal overlap of the contacts of the PS/PI and the hydrophobic region. This suggests that the hydrophobic region is selectively engaged in the side-binding orientation. Indeed, during side-binding, the two PIP binding regions and the hydrophobic region bind simultaneously. When looking closer at the initiation, first the electrostatic interaction occurs followed by the hydrophobic interaction. Furthermore, the lack of membrane binding in the pure POPC system suggests that the electrostatic interaction is crucial for membrane binding.

In summary, our coarse-grained simulations unraveled three distinct binding regions of the C2C domain, one specific for PIP lipids, one for simply negatively charged lipids such as PS and PI, and one for generic lipid, specifically hydrophobic, interactions. The results of our analysis also suggest that the side-binding orientation is result from the PIP and hydrophobic regions interacting co-operatively with the membrane, while the top-binding orientation stems solely from the PS/PI site interactions. The regions are presented graphically in Figure 9.

**Figure 9:**
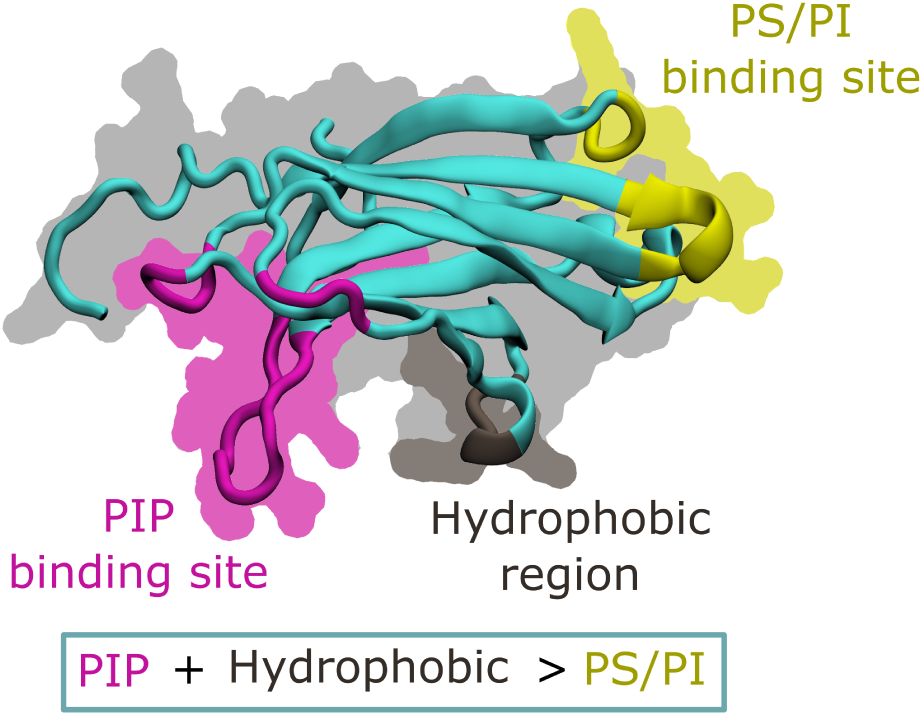
Summary of the lipid-specific binding regions identified in the CG simulations.

### The PS/PI binding site gains impact under PI(4,5)P_2_ break-down

E-Syts specifically operate under conditions of PI(4,5)P_2_ depletion during PLC signaling. We therefore investigated the effect of the PS/PI binding site on the PM localization under conditions of PI(4,5)P_2_ and PI4P break-down. To this end, we expressed the two double mutants R763A K870A and R768A R769A together with the recruitable phosphatase system (Figure 3a). To induce a robust depletion of both PI(4,5)P_2_ and PI4P, we expressed pseudojanin, the combined 4- and 5-phosphatase. Application of rapamycin induced a comparable recruitment of pseudojanin under all conditions (Figure 10c) and induced a strong dissociation of the wild-type C2C domain (Figure 10a,b). Interestingly, the dissociation of the PS/PI binding site mutants was slightly and – in case of the R763A K870A mutant – significantly increased compared to the wild-type domain. This indicates that under conditions of break-down of PI(4,5)P_2_ and PI4P, the PS/PI binding site contributes to sustaining PM localization of the C2C domain. Note that in this experimental set-up, we cannot discriminate between PS and PI binding as both lipids are present during pseudojanin recruitment: PS levels are generally high in the inner leaflet of the PM and PI was generated from PI(4,5)P_2_ and PI4P through dephosphorylation.

**Figure 10:**
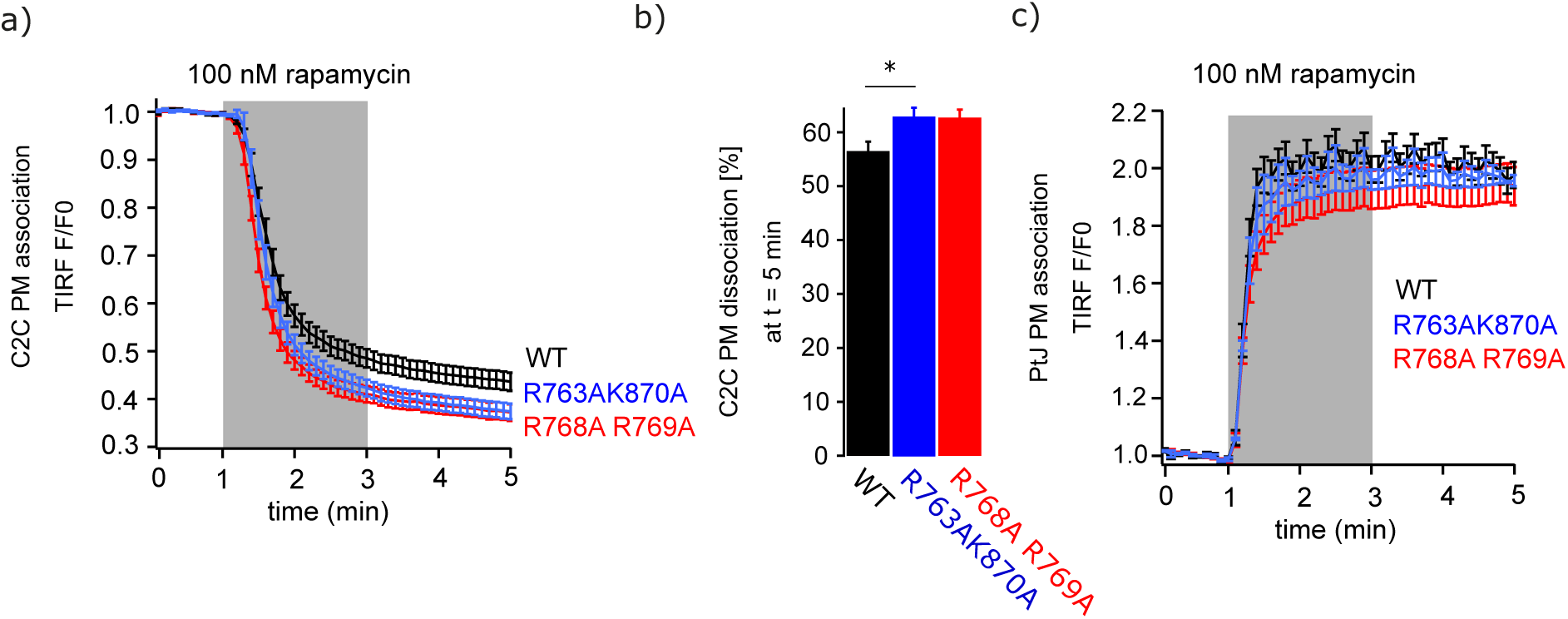
PS/PI binding site slightly contributes to PM association during conditions of phosphoinositide break-down. TIRF imaging of CHO cells expressing GFP-tagged wild-type (WT) C2C and mutant domains, PM-anchored FRB, and RFP-FKBP-pseudojanin. Application of rapamycin induced recruitment of pseudojanin to the PM (c) followed by PI4P and PI(4,5)P_2_ break-down. a) Normalized GFP fluorescence following application of rapamycin. Mean +/-SEM of n cells are shown. WT: n = 51; R763A K870A: n = 36; R768A R769A: n = 40. b) Statistical analysis from experiments in a) with *: p *<* 0.05. c) Normalized RFP fluorescences of the same cells shown in a) confirming successful and comparable PM recruitment of the phosphatases.

## Discussion

### Identification of previously unknown lipid interactions and membrane binding mechanism

With a complementary approach combining in silico and cell biological methods, we identified previously unknown protein-lipid interaction of the C2C domain of E-Syt3 of physiological relevance. Its PM localization depends not only on binding to PI(4,5)P_2_, but also on PI4P binding. Under conditions of phosphoinositide break-down, PS/PI binding might additionally maintain C2C’s PM association.

Further, our computational experiments unraveled the mechanism of membrane association of the C2C domain. In total, we identified three membrane interacting regions on the surface of the C2C domain: a PS/PI binding site, a PIP binding site, and a hydrophobic region. These interaction surfaces allow binding to the membrane either in a top binding mode via the PS/PI binding site or in a side binding mode via the PIP site and hydrophobic region. We show that membrane association is initiated electrostatically via interaction with negatively charged headgroups, but for sustained PM binding the integration of the hydrophobic stretch into the bilayer is needed. Because this hydrophobic interaction occurs only in the side binding configuration, this binding mode accounts for stronger and more sustained membrane association. We confirmed the PIP and PS/PI binding sites in our cellular experiments by means of mutations and identified R813, K814, and R854 as the amino acids most relevant for PM association.

### Comparison of computational and experimental results

Running a computational assay with different membrane compositions, we identified two binding sites for lipid head-groups, namely one for PIPs and another one for PS/PI. Mutation experiments in live cells confirmed that these binding sites mediate membrane binding of the E-Syt3 C2C domain. While the PIP binding site is essential for membrane binding and mutations affect membrane localization under resting conditions (see Figure 7), the PS/PI binding site impacts membrane localization only significantly, when PIP lipids are depleted (see Figure 10).

In our coarse-grained simulations, the impact of the PS/PI binding site seems to be over-estimated in relation to the PIP binding site, when comparing the in silico data with our live-cell experiments. Similarly, the relative binding of PI(4,5)P_2_ and PI(4)P differ in experiment and simulations. These differences might be caused by the different membrane environments: while we used a simple five-component model membrane of the PM’s inner leaflet for the in silico membrane binding assay, the live-cell experiments include the full complexity of the PM. This does not only concern lipid species, but also proteins being present in the plasma membrane as well as the composition of the cytosol. Regarding the relative binding of PI(4,5)P_2_ and PI(4)P, the choice to use a charge of *−*4 for PI(4,5)P_2_ compared to *−*5 corresponding to the fully deprotonated species might cause the difference.^41^ When interacting with the PIP binding site, the pK_a_ of PI(4,5)P_2_ might change leading to full deprotonation. Nevertheless, we could experimentally confirm that the PS/PI binding site impacts membrane binding under conditions of PIP depletion. This highlights the outstanding performance of the Martini 3 force field in identifying protein-lipid interactions.

Moreover, the CG simulations provide insights on the submolecular level which is not accessible experimentally but complements the experimental findings. Based on the simulations, we obtained the molecular binding mechanism of the E-Syt3 C2C domain which shows that the PIP and PS/PI binding sites lead to different binding orientations. Moreover, the PIP binding site is further strengthened by a hydrophobic binding site where W803 inserts in the membrane. This is only possible when the C2C domain binds the membrane with the PIP binding site (see Figure 9).

### Lipid composition-dependent binding orientation

The lipid-specific binding sites result in a lipid composition-dependent binding behavior of the C2C domain of E-Syt3. While this behavior has not been described for the C-terminal C2 domains of E-Syts, there are a number of examples in the literature where other C2 domains exhibit several binding conformations. CG simulations revealed that six Ca^2+^-independent C2 domains from different proteins have individual binding orientation distributions, which additionally depend on the lipid composition.^28^ In the study, three bilayer compositions – namely PC, PC:PS, and PC:PS:PIP_2_ were used. Interestingly, all six tested C2 domains showed already interaction with the zwitterionic PC headgroups, which is in contrast to the E-Syt3 C2C domain. One possible explanation for the difference could be the use of different Martini versions, Martini 2.2 versus Martini 3.0 here. It is known that Martini 2 tends to overestimate interactions.^19,42^

Atomistic MD simulations revealed that the Ca^2+^-dependent C2 domain of protein kinase Cα exhibits lipid-dependent membrane binding as well as binding orientation. ^43^ For membrane binding, negatively charged lipids are required and interestingly, the presence of diacylglycerol increases the width of the binding angle distribution, thus increasing the orientational flexibility of the C2 domain. In line with our results, a pure POPC membrane was not sufficient for membrane binding. A combination of electron paramagnetic resonance measurements and atomistic MD simulations uncovered membrane binding of the C2A domains of synaptotagmin 1 and 7.^44,45^ Similar to our observations, two orientations – one parallel and one perpendicular to the membrane – were present as well as two phenylalanine residues which inserted into the membrane stabilizing the bound state. Also for the C2A domain of dysferlin, a protein involved in membrane resealing, different orientations based on the membrane composition were observed by means of non-linear surface-specific vibrational spectroscopy.^46^ For C2 domains of synaptotagmin 1, Ca^2+^-dependent binding orientations are reported.^47^

### Functional impact of the newly identified lipid interactions

E-Syts are ER-PM tethers and one of their main functions is the dynamic formation and regulation of ER-PM contact sites. Step-wise binding of their multiple C2 domains has been suggested to adjust the ER-PM distance,^48^ which serves as a mechanism to control lipid transfer across ER-PM contact sites.^7,12^

In case of E-Syt3, ER-PM tethering is regulated via PM binding of the C2C domain.^5^ It is Ca^2+^ independent and dynamic tethering is likely regulated via the lipid composition and lipid dynamics at the PM face of the contact sites. Understanding the exact lipid binding properties is fundamental to understand the formation of ER-PM contact sites and regulation of lipid transfer.

Here, we identified that the C2C domain of E-Syt3 binds strongly to PI4P and to a minor extent to PS and PI lipids in addition to its known PI(4,5)P_2_ binding. Under resting conditions, PI4P and PI(4,5)P_2_ levels at the PM are high and both lipids substantially contribute to ER-PM tethering by E-Syt3. However, PI4P binding – and to a certain extend also PS/PI binding – likely gains even more importance during conditions of PI(4,5)P_2_ break-down, e.g. during activation of phospholipase Cβ. Under these conditions, integrity of ER-PM contact sites is needed most in order to supply PI from the ER to the PM and to refill PM phosphoinositide pools. In the past, it was a conundrum as to how E-Syts maintain the integrity of ER-PM contact sites when their PM binding partner PI(4,5)P_2_ is depleted. We suggest that binding of the C2C domain to PI4P helps maintaining the integrity of ER-PM contact sites during PI(4,5)P_2_ break-down. Additionally, binding to PS and/or PI potentially contributes to ER-PM integrity.

In our cellular experiments, the PI(4,5)P_2_ affinity clearly exceeded that of PI4P, as conversion of PI(4,5)P_2_ into PI4P by Inp54 induced a strong dissociation of the C2C domain from the PM. This suggests that PI4P binding cannot fully replace PI(4,5)P_2_ binding during PI(4,5)P_2_ break-down. However, in all our cellular experiments, we investigated the lipid binding properties of the C2C domain in the bulk PM. However, local lipid dynamics at ER-PM contact sites (e.g. during PLCβ activation) might differ from the dynamics in the overall PM. Especially, PI and phophoinositides generated from PI (e.g. PI4P) might accumulate at the PM site of the contact sites during PLCβ activation. We hypothesize that such a local accumulation further increases the relevance of the C2C-PI4P interaction in the integrity of ER-PM contact sites.

From our cellular experiments, we cannot discriminate between PS and PI binding, but as PS levels are generally high at the PM and PI levels are low,^49,50^ we hypothesize that during physiological conditions, PS binding plays a more important role than PI binding. However, as discussed before, the local lipid composition at ER-PM contact sites might differ from that of the general PM also during resting conditions. Especially, PI levels might be locally increased in these sites, because PI transfer occurs in E-Syt-mediated ER-PM contact sites. Such local lipid compositions might shift the physiological relevance of the newly identified C2C-lipid interactions.

We restricted our study to the C2C domain of E-Syt3 which is homologous to the C2C domain of E-Syt2 and the C2E domain of E-Syt1.^10^ Their membrane binding will be part of future investigations to elucidate whether the observed lipid versatility represents a common mechanism within the C2 domains of the E-Syt family. For E-Syt2, a basic patch has already been identified to contribute to PM localization under resting conditions.^5^

## Conclusions

In summary, we identified three previously unknown lipid binding sites of the C2C domain of E-Syt3 by combining coarse-grained MD simulations and live-cell mutation experiments. These binding sites exhibit different lipid specificities, namely preferring PIP lipids, PS/PI lipids as well as a hydrophobic region which inserts into the hydrophobic membrane core. The following molecular picture emerges from the CG simulations: the PIP site and hydrophobic region act together and support strong membrane binding. The PS/PI site induces a different orientation of the C2C domain on the membrane surface and does not allow for hydrophobic binding.

In live cells, the strongest binding occurs via the PIP site. The PS/PI site only comes into play when PIP lipids are depleted. Therefore we hypothesize that the PS/PI site acts as a safeguard mechanism to maintain integrity of ER-PM contact sites during conditions of PI(4,5)P_2_ break-down in the inner leaflet of the PM.

In the future, we will study further domains of the extended synaptotagmin protein family to obtain a better understanding of the role of different lipid binding sites for regulation and stability of E-Syt-mediated ER-PM contact sites.

## Materials and Methods

### Computational Methods

All simulation systems were modeled with the Martini 3 force field^19,51^ and the GōMartini 3 model was used for the C2C domain.^52^ The protein structure for E-Syt3 (Uniprot:A0FGR9) was retrieved from Alphafold2 structure prediction database (AF-A0FGR9-F1-v4).^53–55^ The C2C domain residues from 745-886 were extracted from the full structure using PyMol.^56^ Next, overlap (OV) and repulsive contacts of structural units (rCSU) contact maps^57^ were calculated for the model using the Gō Contact Map server^58^ with the default settings (sequence distance=4, cut-off_long_ = 1.1 nm and cut-off_short_ = 0.3 nm). The protein structure was then coarse-grained with Martinize2^59^ with the atomistic side-chain orientation correction^60^ and automated disulfide bridge detection. The secondary structure for the domain was determined using DSSPs 3.1.4,^61,62^ and the secondary and tertiary structures were set with 281 Gō-like bonds with Lennart-Jones potential of *ɛ_LJ_* = 12.0 kJ/mol.

The simulation systems with lipid bilayers were set up using the insane.py script^41,63^ in a neutral box with physiological 0.15 M NaCl concentration. In all setups the C2C domain was positioned in the water phase approximately 2 nm above the membrane in the simulation box, in order to start the simulation in an unbiased fully solvated conformation. The exact composition of the membrane as well as the solvent are given in Supplementary Table S1 in the SI; exact sizes of simulation boxes with particle counts for each system in Supplementary Table S2 in the SI. Three replicas of each system were run for at least 10 µs with GROMACS 2020.5.^64^ The simulations were run using a timestep of 20 fs, with the temperature set to 310 K with V-rescale temperature coupling using a coupling constant of 1 ps; the pressure of 1 bar was set with a semi-isotropic Parrinello-Rahman barostat using a coupling constant of 12 ps and compressibility of 3 *·* 10*^−^*^4^ bar*^−^*^1^. Electrostatic and van-der-Waals interactions were handled with the new reaction-field approach, with a reaction field potential of *ɛ_rf_* = 15 within a cutoff of 1.1 nm and set to infinity beyond the cutoff.^65^ To avoid the unwanted neighbour-list artifacts, the verlet buffer tolerance was set to 0.0002 kJmol*^−^*^1^ps*^−^*^1^ and the neighbour list energy, temperature coupling, and pressure coupling were set to be calculated every 20 steps.^66^ To facilitate better sampling of the phase-space, random starting velocities were created for the starting structures of each of the two other repeats.

### Analysis

The final trajectories were analysed using a combination of GROMACS analysis tools and in-house Python scripts.

The distance between the C2C domain and the PO4 layer of the membrane was determined to be the z-distance between the center of geometry (COG) of the C2C domain and the PO4 bead plane of the closest leaflet. The analysis was conducted by using gmx distance with option -xyz set between the COG of the C2C domain plus the COG of PO4 beads in either the upper or lower leaflet. Due to the symmetry of the system and periodic boundary conditions, the shortest z-distance of the two measurements was selected as the final result for each snapshot. The distance distributions were calculated from the combined data of all three repeats to bargraphs with 80 bins of a bin width of approximately 0.08 nm. The histogram was normalized to 1 to ensure equal scale across the simulation sets regardless of the exact length of the individual simulations.

The analysis of the orientation angle of the protein was conducted using gmx gangle by determining the smallest angle between the protein orientation vector and the membrane normal. The protein orientation vector was defined from the backbone bead of residue threonine 782 (T782) to the backbone bead of methionine 820 (M820), and spanned approximately the principal axis of the C2C-domain. For the angle distributions, the data first needed to be corrected for equatorial sampling bias to ensure uniform distribution of angles in the unbound state. The corrected distributions were then determined similarly to the distance distributions with normalization to 1. To create the 2D orientation-distance histograms, the data from distance analysis and angle analysis were paired for each snapshot, the angle columns were corrected for equatorial sampling bias, the total area of the histogram was normalized to 1, and finally, the distribution was plotted as 2D histogram. The distributions and normalizations were completed using an in-house Python script. The contact analysis was conducted using gmx select for the selected group of beads (*e.g.* whole protein, residue, or group of residues) as a function of time. A lipid molecule was counted to be in contact with the protein or an individual residue if it was within 0.6 nm of the protein/residue beads. If multiple beads in one lipid molecule were within that distance, they were counted as one contact regardless. The average number of contacts per residue was calculated as the combined average over all repeats of the system.

### Cell culture

CHO dhFr-cells were cultured in MEM Alpha medium (gibco, ThermoFisher Scientific, Waltham, US), supplemented with 10% fetal calf serum, 1% penicillin and 1% streptomycin. Cells were grown at 37°C and 5 % CO2. Cells were seeded on cover slips for confocal imaging or on glass bottom µ-slide VI0.5 flow chambers (ibidi, Martinsried, Germany) for TIRF imaging. Transfections were performed 24h pre imaging using polyethylenimine (PEI).

### Molecular biology

Expression constructs used for transfection were: Lyn11-FRB (GCIKSKGKDSA and AA 2026-2121 of FRB (NM 004958.3)) in pECFP-N1^40^ and in pC4-RHE;mCherry-FKBP-pseudojanin in pEGFP-C1 (AA 30-108 of FKBP, NM 054014.3; AA 2-517 of SAC1, NM 001179777.1; AA 214-644 of InPP5E, NM 019892.4;^39,40^ CFP-FKBP-Inp54p (AA 30-108 of FKBP, NM 054014.3; Inp54p: Z74807,^67^ L10-Anker in pRFP, C2C domain of human E-Syt3 in pRFP-C1 (AA 745-887; NM 031913.4^5^). For the generation of the RFP-tagged C2C domain of E-Syt3, the E-Syt3 portion was amplified by PCR and cloned into pRFP-C3 through HindIII and KpnI restriction sites. The GFP-tagged construct was generated via fluorophore exchange using BshT1 and Bsp1407I restriction sites. Point mutations were introduced into the GFP-tagged construct via site-directed mutagenesis.

### Confocal microscopy

Confocal imaging was done on a Zeiss Examiner upright microscope equipped with a LSM710 scan head (Carl Zeiss AG, Jena, Germany) and a W-Plan-Apo-chromat 20x/1.0 DIC M27 75 mm (Carl Zeiss AG). For RFP excitation a 561 laser line and for GFP a 488 laser line was used. Emission filters ranged from 620/62 for RFP and from 525/50 for GFP. Image acquisition was done with ZEN software (Carl Zeiss AG). During imaging cells were kept in extracellular solution (5.8 mM KCl, 144 mM NaCl, 0.9 mM MgCl2, 1.3 mM CaCl2, 0.7 mM NaH2PO4, 5.6 mM D-glucose, 10 mM HEPES, pH = 7.4).

### TIRF microscopy

TIRF experiments were performed as described in reference.^33^ In brief, imaging was performed on a Dmi8 upright microscope (Leica, Wetzlar, Germany) equipped with an Infinity TIRF module (Leica), a HC PL APO 100x/1.47 OIL objective (Leica) and a widefield laser (Leica). GFP-, RFP- and CFP-fluorescence was excited at 488 nm, 561 nm and 405 nm, respectively. During imaging cells were superfused with extracellular solution (ECS) (5.8 mM KCl, 144 mM NaCl, 0.9 mM MgCl2, 1.3 mM CaCl2, 0.7 mM NaH2PO4, 5.6 mM D-glucose, 10 mM HEPES, pH = 7.4). Rapamycin was purchased from Calbiochem (Merck Chemicals GmbH, Darmstadt, Germany) as 2.5 mM stock solution in DMSO and diluted in extracellular solution to a working solution of 1 µM. Rapamycin was applied as indicated in the figure legends, images were taken every 6 s.

### Imaging data analysis

Imaging data was analyzed with ImageJ software and IGOR Pro (WaveMetrics, Lake Oswego, OR, USA). Statistical evaluation was done with IGOR Pro. Data are displayed as mean ± SEM of individual cells. Statistical tests were done on individual cells. For multi-comparisons Scheffe tests were performed. Asterisks indicate significance levels: *: p *<* 0.05; **: p *<* 0.01; ***: p *<* 0.001.

## Acknowledgement

S.L. and S.T. gratefully acknowledge the computing time provided at the NHR Center NHR@SW at Goethe University Frankfurt (project number 26814). This is funded by the Federal Ministry of Education and Research, and the state governments participating on the basis of the resolutions of the GWK for national high performance computing at universities (www.nhr-verein.de/unsere-partner). This work was funded by the Deutsche Forschungsgemeinschaft (DFG, German Research Foundation) under Germany’s Excellence Strategy – EXC-3094 – 533751785 (to F.W. and S.T.), the Hessian Ministry of Science and Arts LOEWE program (Center for Multiscale Modeling in Life Sciences, CMMS) (to S.T.), Alfons und Gertrud Kassel foundation (to S.T.), and Dr. Rolf M. Schwiete foundation (to S.T.).

## Supporting Information Available

Supporting Tables and Figures are available.

## Author Information

### Contributions

V.T. and S.T. initiated the project and designed the research. S.L. performed and analyzed the MD simulations. V.T. performed and analyzed the experiments with help of T.J.. All authors interpreted the data and discussed the results. S.L., V.T., and S.T. wrote the manuscript which was reviewed by all authors. F.W., D.O., and S.T. provided funding. V.T. and S.T. supervised the research and coordinated the project.

## References

(1) Okeke, E.; Dingsdale, H.; Parker, T.; Voronina, S.; Tepikin, A. V. Endoplasmic reticulum–plasma membrane junctions: structure, function and dynamics. The Journal of Physiology 2016, 594, 2837–2847, eprint: https://physoc.onlinelibrary.wiley.com/doi/pdf/10.1113/JP271142.

(2) Prinz, W. A.; Toulmay, A.; Balla, T. The functional universe of membrane contact sites. Nat Rev Mol Cell Biol 2020, 21, 7–24, Number: 1.

(3) Scorrano, L.; De Matteis, M. A.; Emr, S.; Giordano, F.; Hajnoczky, G.; Kornmann, B.; Lackner, L. L.; Levine, T. P.; Pellegrini, L.; Reinisch, K.; Rizzuto, R.; Simmen, T.; Stenmark, H.; Ungermann, C.; Schuldiner, M. Coming together to define membrane contact sites. Nat Commun 2019, 10, 1287, Number: 1.

(4) Kors, S.; Kurian, S. M.; Costello, J. L.; Schrader, M. Controlling contacts—Molecular mechanisms to regulate organelle membrane tethering. BioEssays 2022, 44, 2200151, eprint: https://onlinelibrary.wiley.com/doi/pdf/10.1002/bies.202200151.

(5) Giordano, F.; Saheki, Y.; Idevall-Hagren, O.; Colombo, S. F.; Pirruccello, M.; Milosevic, I.; Gracheva, E. O.; Bagriantsev, S. N.; Borgese, N.; De Camilli, P. PI(4,5)P2-Dependent and Ca2+-Regulated ER-PM Interactions Mediated by the Extended Synaptotagmins. Cell 2013, 153, 1494–1509.

(6) Schauder, C. M.; Wu, X.; Saheki, Y.; Narayanaswamy, P.; Torta, F.; Wenk, M. R.; De Camilli, P.; Reinisch, K. M. Structure of a lipid-bound extended synaptotagmin indicates a role in lipid transfer. Nature 2014, 510, 552–555.

(7) Saheki, Y.; Bian, X.; Schauder, C. M.; Sawaki, Y.; Surma, M. A.; Klose, C.; Pincet, F.; Reinisch, K. M.; De Camilli, P. Control of plasma membrane lipid homeostasis by the extended synaptotagmins. Nat Cell Biol 2016, 18, 504–515.

(8) Wang, Y.; Li, Z.; Wang, X.; Zhao, Z.; Jiao, L.; Liu, R.; Wang, K.; Ma, R.; Yang, Y.; Chen, G.; Wang, Y.; Bian, X. Insights into membrane association of the SMP domain of extended synaptotagmin. Nat Commun 2023, 14, 1504, Number: 1.

(9) Idevall-Hagren, O.; Lu, A.; Xie, B.; De Camilli, P. Triggered Ca2+ influx is required for extended synaptotagmin 1-induced ER-plasma membrane tethering. The EMBO Journal 2015, 34, 2291–2305, Num Pages: 2305.

(10) Min, S.-W.; Chang, W.-P.; Sudhof, T. C. E-Syts, a family of membranous Ca2+-sensor proteins with multiple C2 domains. Proceedings of the National Academy of Sciences 2007, 104, 3823–3828.

(11) Bian, X.; Saheki, Y.; De Camilli, P. Ca2+ releases E-Syt1 autoinhibition to couple ER-plasma membrane tethering with lipid transport. EMBO J 2017, 37, 6.

(12) Chang, C.-L.; Hsieh, T.-S.; Yang, T. T.; Rothberg, K. G.; Azizoglu, D. B.; Volk, E.; Liao, J.-C.; Liou, J. Feedback Regulation of Receptor-Induced Ca2+ Signaling Mediated by E-Syt1 and Nir2 at Endoplasmic Reticulum-Plasma Membrane Junctions. Cell Reports 2013, 5, 813–825.

(13) Chang, C.-L.; Liou, J. Phosphatidylinositol 4,5-Bisphosphate Homeostasis Regulated by Nir2 and Nir3 Proteins at Endoplasmic Reticulum-Plasma Membrane Junctions *. Journal of Biological Chemistry 2015, 290, 14289–14301.

(14) Balla, T. Phosphoinositides: Tiny Lipids With Giant Impact on Cell Regulation. Physiological Reviews 2013, 93, 1019–1137.

(15) Kim, S.; Kedan, A.; Marom, M.; Gavert, N.; Keinan, O.; Selitrennik, M.; Laufman, O.; Lev, S. The phosphatidylinositol-transfer protein Nir2 binds phosphatidic acid and positively regulates phosphoinositide signalling. EMBO Rep 2013, 14, 13.

(16) Kim, Y. J.; Guzman-Hernandez, M.-L.; Wisniewski, E.; Balla, T. Phosphatidylinositol-Phosphatidic Acid Exchange by Nir2 at ER-PM Contact Sites Maintains Phosphoinositide Signaling Competence. Developmental Cell 2015, 33, 549–561.

(17) Fernández-Busnadiego, R.; Saheki, Y.; De Camilli, P. Three-dimensional architecture of extended synaptotagmin-mediated endoplasmic reticulum–plasma membrane contact sites. Proceedings of the National Academy of Sciences 2015, 112, E2004–E2013.

(18) Corradi, V.; Sejdiu, B. I.; Mesa-Galloso, H.; Abdizadeh, H.; Noskov, S. Y.; Marrink, S. J.; Tieleman, D. P. Emerging Diversity In Lipid-Protein Interactions. Chemical reviews 2018, 119, 5775–5848.

(19) Souza, P. C. T. et al. Martini 3: a general purpose force field for coarse-grained molecular dynamics. Nature Methods 2021, 18, 382–388.

(20) Ingolfsson, H. I.; Melo, M. N.; van Eerden, F. J.; Arnarez, C.; Lopez, C. A.; Wassenaar, T. A.; Periole, X.; de Vries, A. H.; Tieleman, D. P.; Marrink, S. J. Lipid Organization of the Plasma Membrane. Journal of the American Chemical Society 2014, 136, 14554–14559, PMID: 25229711.

(21) Pedersen, K. B. et al. The Martini 3 Lipidome: Expanded and Refined Parameters Improve Lipid Phase Behavior. ACS Cent. Sci. 2025, 11, 1598–1610.

(22) Borges-Araujo, L.; Borges-Araujo, A. C.; Ozturk, T. N.; Ramirez-Echemendia, D. P.; Fabian, B.; Carpenter, T. S.; Thallmair, S.; Barnoud, J.; Ingolfsson, H. I.; Hummer, G.; Tieleman, D. P.; Marrink, S. J.; Souza, P. C. T.; Melo, M. N. Martini 3 Coarse-Grained Force Field for Cholesterol. Journal of Chemical Theory and Computation 2023, 19, 7387–7404.

(23) Corradi, V.; Bukiya, A. N.; Miranda, W. E.; Cui, M.; Plant, L. D.; Logothetis, D. E.; Tieleman, D. P.; Noskov, S. Y.; Rosenhouse-Dantsker, A. A molecular switch controls the impact of cholesterol on a Kir channel. Proceedings of the National Academy of Sciences 2022, 119, e2109431119.

(24) Lin, Y.; Tao, E.; Champion, J. P.; Corry, B. A binding site for phosphoinositides described by multiscale simulations explains their modulation of voltage-gated sodium channels. eLife 2024, 12, RP91218.

(25) Arnarez, C.; Mazat, J.-P.; Elezgaray, J.; Marrink, S.-J.; Periole, X. Evidence for Cardiolipin Binding Sites on the Membrane-Exposed Surface of the Cytochrome *bc* _1_. Journal of the American Chemical Society 2013, 135, 3112–3120.

(26) Corey, R. A.; Song, W.; Duncan, A. L.; Ansell, T. B.; Sansom, M. S. P.; Stansfeld, P. J. Identification and assessment of cardiolipin interactions with E. coli inner membrane proteins. Science Advances 2021, 7, eabh2217.

(27) Ansell, T. B.; Song, W.; Coupland, C. E.; Carrique, L.; Corey, R. A.; Duncan, A. L.; Cassidy, C. K.; Geurts, M. M. G.; Rasmussen, T.; Ward, A. B.; Siebold, C.; Stansfeld, P. J.; Sansom, M. S. P. LipIDens: simulation assisted interpretation of lipid densities in cryo-EM structures of membrane proteins. Nat Commun 2023, 14, 7774.

(28) Larsen, A. H.; Sansom, M. S. P. Binding of Ca2+-independent C2 domains to lipid membranes: A multi-scale molecular dynamics study. *Structure (London*, England : 1993) 2021, 29, 1200–1213.e2.

(29) John, L. H.; Naughton, F. B.; Sansom, M. S. P.; Larsen, A. H. The Role of C2 Domains in Two Different Phosphatases: PTEN and SHIP2. Membranes 2023, 13, 408, Number: 4.

(30) Yamamoto, E.; Domanski, J.; Naughton, F. B.; Best, R. B.; Kalli, A. C.; Stansfeld, P. J.; Sansom, M. S. P. Multiple lipid binding sites determine the affinity of PH domains for phosphoinositide-containing membranes. Science Advances 2020, 6, eaay5736.

(31) Naughton, F. B.; Kalli, A. C.; Sansom, M. S. P. Association of Peripheral Membrane Proteins with Membranes: Free Energy of Binding of GRP1 PH Domain with Phosphatidylinositol Phosphate-Containing Model Bilayers. The Journal of Physical Chemistry Letters 2016, 7, 1219–1224.

(32) Thallmair, V.; Schultz, L.; Zhao, W.; Marrink, S. J.; Oliver, D.; Thallmair, S. Two cooperative binding sites sensitize PI(4,5)P2 recognition by the tubby domain. Science Advances 2022, 8, eabp9471.

(33) Thallmair, V.; Schultz, L.; Evers, S.; Jolie, T.; Goecke, C.; Leitner, M. G.; Thallmair, S.; Oliver, D. Localization of the tubby domain, a PI(4,5)P2 biosensor, to E-Syt3-rich endoplasmic reticulum–plasma membrane junctions. Journal of Cell Science 2023, 136, jcs260848.

(34) Khan, H. M.; Souza, P. C. T.; Thallmair, S.; Barnoud, J.; de Vries, A. H.; Marrink, S. J.; Reuter, N. Capturing Choline–Aromatics Cationπ Interactions in the MARTINI Force Field. J. Chem. Theory Comput. 2020, 16, 2550–2560.

(35) Cheung, L. K. Y.; Thallmair, S.; Yada, R. Y. Elucidating the structure and function of a membrane-active plant protein domain using in silico mutagenesis. Biochimica et Biophysica Acta (BBA) - Biomembranes 2025, 1867, 184409.

(36) Ingolfsson, H. I.; Bhatia, H.; Zeppelin, T.; Bennett, W. F. D.; Carpenter, K. A.; Hsu, P.-C.; Dharuman, G.; Bremer, P.-T.; Schiøtt, B.; Lightstone, F. C.; Carpenter, T. S. Capturing Biologically Complex Tissue-Specific Membranes at Different Levels of Compositional Complexity. The Journal of Physical Chemistry B 2020, 124, 7819–7829, PMID: 32790367.

(37) Horvath, S. E.; Daum, G. Lipids of mitochondria. Progress in Lipid Research 2013, 52, 590–614.

(38) Pogozheva, I. D.; Armstrong, G. A.; Kong, L.; Hartnagel, T. J.; Carpino, C. A.; Gee, S. E.; Picarello, D. M.; Rubin, A. S.; Lee, J.; Park, S.; Lomize, A. L.; Im, W. Comparative Molecular Dynamics Simulation Studies of Realistic Eukaryotic, Prokaryotic, and Archaeal Membranes. Journal of Chemical Information and Modeling 2022, 62, 1036–1051.

(39) Lindner, M.; Leitner, M. G.; Halaszovich, C. R.; Hammond, G. R. V.; Oliver, D. Probing the regulation of TASK potassium channels by PI(4,5)P2 with switchable phosphoinositide phosphatases. The Journal of Physiology 2011, 589, 3149–3162.

(40) Hammond, G. R. V.; Fischer, M. J.; Anderson, K. E.; Holdich, J.; Koteci, A.; Balla, T.; Irvine, R. F. PI4P and PI(4,5)P2 Are Essential But Independent Lipid Determinants of Membrane Identity. Science 2012, 337, 727–730.

(41) Borges-Araujo, L.; Souza, P. C. T.; Fernandes, F.; Melo, M. N. Improved Parameterization of Phosphatidylinositide Lipid Headgroups for the Martini 3 Coarse-Grain Force Field. Journal of Chemical Theory and Computation 2022, 18, 357–373, PMID: 34962393.

(42) Alessandri, R.; Souza, P. C. T.; Thallmair, S.; Melo, M. N.; de Vries, A. H.; Marrink, S. J. Pitfalls of the Martini Model. Journal of Chemical Theory and Computation 2019, 15, 5448–5460.

(43) Alwarawrah, M.; Wereszczynski, J. Investigation of the Effect of Bilayer Composition on PKCα-C2 Domain Docking Using Molecular Dynamics Simulations. The Journal of Physical Chemistry B 2016, acs.jpcb.6b10188.

(44) Osterberg, J. R.; Chon, N. L.; Boo, A.; Maynard, F. A.; Lin, H.; Knight, J. D. Membrane Docking of the Synaptotagmin 7 C2A Domain: Electron Paramagnetic Resonance Measurements Show Contributions from Two Membrane Binding Loops. Biochemistry 2015, 54, 5684–5695.

(45) Chon, N. L.; Osterberg, J. R.; Henderson, J.; Khan, H. M.; Reuter, N.; Knight, J. D.; Lin, H. Membrane Docking of the Synaptotagmin 7 C2A Domain: Computation Reveals Interplay between Electrostatic and Hydrophobic Contributions. Biochemistry 2015, 54, 5696–5711.

(46) Carpenter, A. P.; Khuu, P.; Weidner, T.; Johnson, C. P.; Roeters, S. J.; Baio, J. E. Orientation of the Dysferlin C2A Domain is Responsive to the Composition of Lipid Membranes. J. Phys. Chem. B 2023, 127, 577–589.

(47) Bender, J.; Kundlacz, T.; Rudden, L. S. P.; Frick, M.; Bieber, J.; Degiacomi, M. T.; Schmidt, C. Ca2+-dependent lipid preferences shape synaptotagmin-1 C2A and C2B dynamics: Insights from experiments and simulations. Structure 2024, 32, 1691– 1704.e5.

(48) Ge, J.; Bian, X.; Ma, L.; Cai, Y.; Li, Y.; Yang, J.; Karatekin, E.; De Camilli, P.; Zhang, Y. Stepwise membrane binding of extended synaptotagmins revealed by optical tweezers. Nat Chem Biol 2022, 18, 313–320, Number: 3.

(49) Zewe, J. P.; Miller, A. M.; Sangappa, S.; Wills, R. C.; Goulden, B. D.; Hammond, G. R. Probing the subcellular distribution of phosphatidylinositol reveals a surprising lack at the plasma membrane. J Cell Biol 2020, 219.

(50) Pemberton, J. G.; Kim, Y. J.; Humpolickova, J.; Eisenreichova, A.; Sengupta, N.; Toth, D. J.; Boura, E.; Balla, T. Defining the subcellular distribution and metabolic channeling of phosphatidylinositol. J Cell Biol 2020, 219.

(51) Marrink, S. J.; Risselada, H. J.; Yefimov, S.; Tieleman, D. P.; Vries, A. H. d. The MARTINI Force Field: Coarse Grained Model for Biomolecular Simulations. The Journal of Physical Chemistry B 2007, 111, 7812–7824.

(52) Souza, P. C. T. et al. GoMartini 3: From large conformational changes in proteins to environmental bias corrections. Nat Commun 2025, 16, 4051.

(53) Database, A. P. S. AlphaFold: AI-driven protein structure predictions. https://alphafold.ebi.ac.uk/, 2021; Accessed: 2024-10-04.

(54) Varadi, M. et al. AlphaFold Protein Structure Database: massively expanding the structural coverage of protein-sequence space with high-accuracy models. Nucleic Acids Research 2021, 50, D439–D444.

(55) Varadi, M. et al. AlphaFold Protein Structure Database in 2024: providing structure coverage for over 214 million protein sequences. Nucleic Acids Research 2023, 52, D368–D375.

(56) PyMOL 2.5.0 Open-Source Molecular Graphics System. Schrodinger, LLC, 2010.

(57) Wolek, K.; Gomez-Sicilia, A.; Cieplak, M. Determination of contact maps in proteins: A combination of structural and chemical approaches. The Journal of Chemical Physics 2015, 143, 243105.

(58) GoContactMap: Analysis and visualization of molecular contact maps. http://pomalab.ippt.pan.pl/GoContactMap/, Accessed: 2024-10-04.

(59) Kroon, P. C.; Grunewald, F.; Barnoud, J.; Tilburg, M. v.; Brasnett, C.; Souza, P. C. T. d.; Wassenaar, T. A.; Marrink, S.-J. J. Martinize2 and Vermouth: Unified Framework for Topology Generation. eLife 2025, 12.

(60) Herzog, F. A.; Braun, L.; Schoen, I.; Vogel, V. Improved Side Chain Dynamics in MARTINI Simulations of Protein–Lipid Interfaces. Journal of Chemical Theory and Computation 2016, 12, 2446–2458.

(61) Kabsch, W.; Sander, C. Dictionary of protein secondary structure: Pattern recognition of hydrogen-bonded and geometrical features. Biopolymers 1983, 22, 2577–2637.

(62) Touw, W. G.; Baakman, C.; Black, J.; te Beek, T. A.; Krieger, E.; Joosten, R. P.; Vriend, G. A series of PDB-related databanks for everyday needs. Nucleic Acids Research 2014, 43, D364–D368.

(63) Wassenaar, T. A.; Ingolfsson, H. I.; Bockmann, R. A.; Tieleman, D. P.; Marrink, S. J. Computational Lipidomics with insane: A Versatile Tool for Generating Custom Membranes for Molecular Simulations. Journal of Chemical Theory and Computation 2015, 11, 2144–2155.

(64) Abraham, M. J.; Murtola, T.; Schulz, R.; Pall, S.; Smith, J. C.; Hess, B.; Lindahl, E. GROMACS: High performance molecular simulations through multi-level parallelism from laptops to supercomputers. SoftwareX 2015, 1, 19–25.

(65) de Jong, D. H.; Baoukina, S.; Ingolfsson, H. I.; Marrink, S. J. Martini straight: Boosting performance using a shorter cutoff and GPUs. Computer Physics Communications 2016, 199, 1–7.

(66) Kim, H.; Fabian, B.; Hummer, G. Neighbor List Artifacts in Molecular Dynamics Simulations. Journal of Chemical Theory and Computation 2023, 19, 8919–8929, PMID: 38035387.

(67) Suh, B.-C.; Inoue, T.; Meyer, T.; Hille, B. Rapid Chemically Induced Changes of PtdIns(4,5)P2 Gate KCNQ Ion Channels. Science 2006, 314, 1454–1457.

