## Supplementary Information for "Lipid versatility of the C2C domain of Extended Synaptotagmin 3 safeguards plasma membrane binding"

Saara Lautala,[†,‡,¶,#] Theresa Jolie,[§] Florian Wilfling,[‡,‖] Dominik Oliver,[§] Veronika Thallmair,[*,§,#] and Sebastian Thallmair[*,†,⊥,‖]

†*Frankfurt Institute for Advanced Studies, Ruth-Moufang-Str. 1, 60438 Frankfurt am Main, Germany*

‡*Mechanisms of Cellular Quality Control, Max-Planck-Institute for Biophysics, 60438 Frankfurt am Main, Germany*

¶*Current affiliation: Schrödinger GmbH, Glücksteinallee 25, 68163 Mannheim, Germany.*

§*Institute for Physiology and Pathophysiology, Philipps University Marburg, Deutschhausstr. 1-2, 35037 Marburg, Germany*

‖*Cluster of Excellence SubCellular Architecture of Life (SCALE), Goethe University Frankfurt, Frankfurt am Main, Germany*

⊥*Faculty of Biochemistry, Chemistry and Pharmacy, Goethe University Frankfurt, 60438 Frankfurt am Main, Germany*

#*These authors contributed equally to this work.*

### Supplementary Tables

Supplementary Table S1: **Composition of the simulated systems.** Lipid composition and solvent particles of all simulated systems. Note that the solvent columns include also counterions as well as 0.15 M NaCl. Each system contained additionally one E-Syt3 C2C domain.

| Membrane model | Lipids | Count | Rel. amount | Solvent | Count |
|---|---|---|---|---|---|
| 5MIX | CHOL | 180 | 0.28 | water | 20,545 |
| | POPC | 162 | 0.25 | $Na^+$ | 423 |
| | POPE | 162 | 0.25 | $Cl^-$ | 222 |
| | POPS | 116 | 0.18 | | |
| | PI(4,5)$P_2$ | 24 | 0.04 | | |
| $5MIX_{doped}$ | CHOL | 174 | 0.27 | water | 19,723 |
| | POPC | 154 | 0.24 | $Na^+$ | 512 |
| | POPE | 154 | 0.24 | $Cl^-$ | 213 |
| | POPS | 110 | 17 | | |
| | PI(4,5)$P_2$ | 50 | 0.08 | | |
| $5MIX_{PI4P}$ | CHOL | 180 | 0.28 | water | 20,545 |
| | POPC | 162 | 0.25 | $Na^+$ | 399 |
| | POPE | 162 | 0.25 | $Cl^-$ | 222 |
| | POPS | 116 | 0.18 | | |
| | PI(4)P | 24 | 0.04 | | |
| $5MIX_{PI}$ | CHOL | 180 | 0.28 | water | 20,545 |
| | POPC | 162 | 0.25 | $Na^+$ | 351 |
| | POPE | 162 | 0.25 | $Cl^-$ | 222 |
| | POPS | 116 | 0.18 | | |
| | PI | 24 | 0.04 | | |
| 4MIX | CHOL | 186 | 0.29 | water | 20,815 |
| | POPC | 168 | 0.26 | $Na^+$ | 336 |
| | POPE | 168 | 0.26 | $Cl^-$ | 225 |
| | POPS | 122 | 0.19 | | |
| POPC | POPC | 644 | 1.00 | water | 25,094 |
| | | | | $Na^+$ | 271 |
| | | | | $Cl^-$ | 282 |

Supplementary Table S2: **System size and simulation time of all systems.** The box size is taken after equilibration of the system.

| Membrane model | Box size ($nm^3$) | Number of particles | Simulation time (µs) |
|---|---|---|---|
| 5MIX | 13.2×13.2×18.2 | 28,975 | 11.3 |
| | | | 11.2 |
| | | | 10.2 |
| $5MIX_{doped}$ | 13.1×13.1×18.0 | 28,357 | 12.1 |
| | | | 12.0 |
| | | | 12.0 |
| $5MIX_{PI4P}$ | 13.2×13.2×18.3 | 28,927 | 10.9 |
| | | | 15.7 |
| | | | 15.2 |
| $5MIX_{PI}$ | 13.2×13.2×18.3 | 28,855 | 10.2 |
| | | | 10.2 |
| | | | 10.3 |
| 4MIX | 13.3×13.3×18.3 | 29,023 | 10.0 |
| | | | 10.3 |
| | | | 10.3 |
| POPC | 13.7×13.7×20.5 | 33,852 | 10.5 |
| | | | 11.3 |
| | | | 11.3 |
| **Total simulation time** | | | **205.0** |

### Supplementary Figures

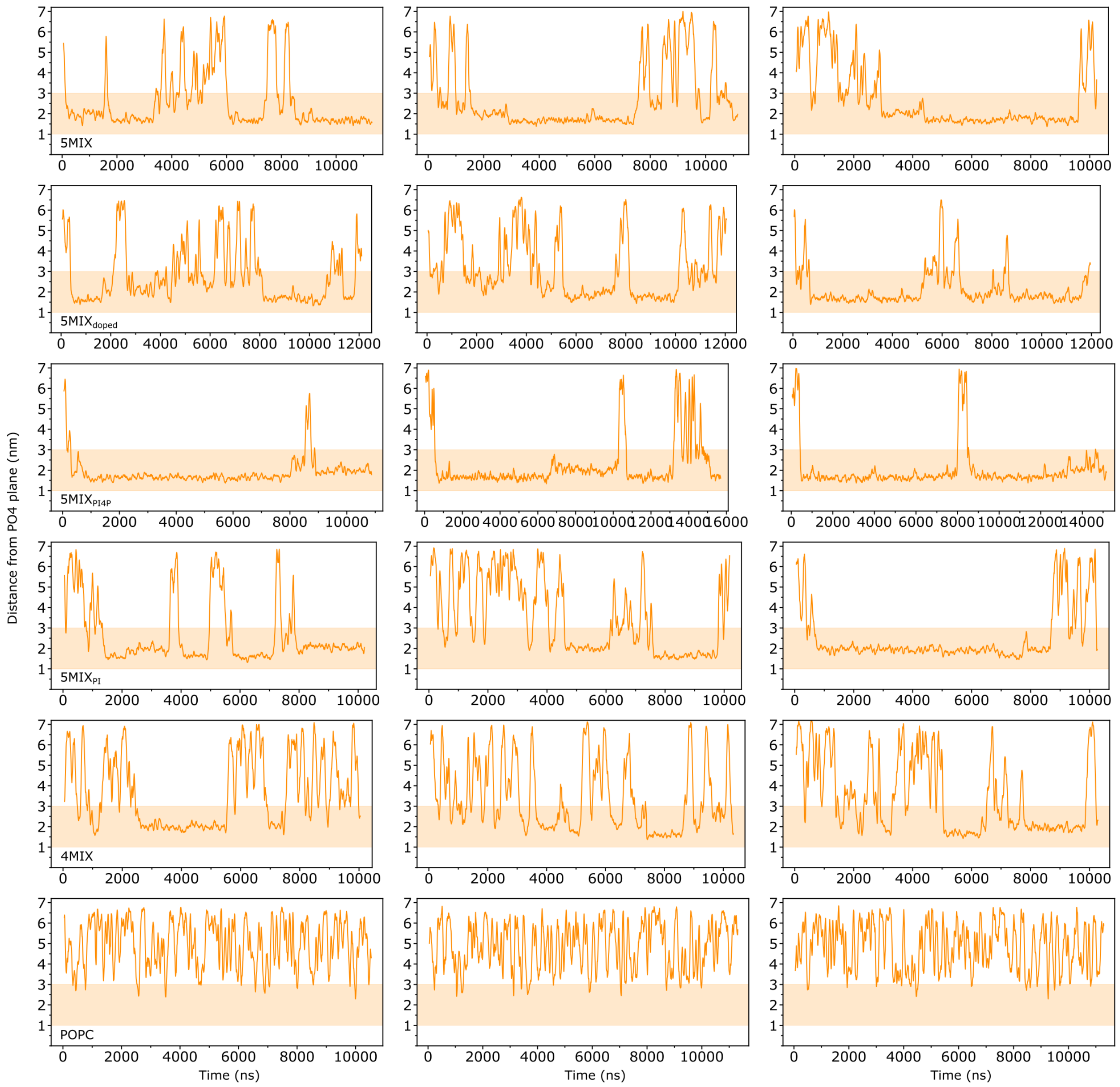


Supplementary Figure S1: **Protein-membrane distance analysis for all replicas.** Plots demonstrating the dynamics of each systems via time-dependent series of the C2C domain's distance to the membrane. The y-axis represents the z-distance between the center of geometry (COG) of the C2C domain and the phosphate bead plane (PO4 plane) of the nearest leaflet. Occupancy inside the highlighted region indicates contact between the C2C domain and the lipid bilayer. Each row shows the three replicas for one lipid composition: 5MIX, 5MIX$_{\mathrm{doped}}$, 5MIX$_{\mathrm{PI4P}}$, 5MIX$_{\mathrm{PI}}$, 4MIX, and POPC. For the exact lipid compositions, see Figure 2a in the main manuscript.

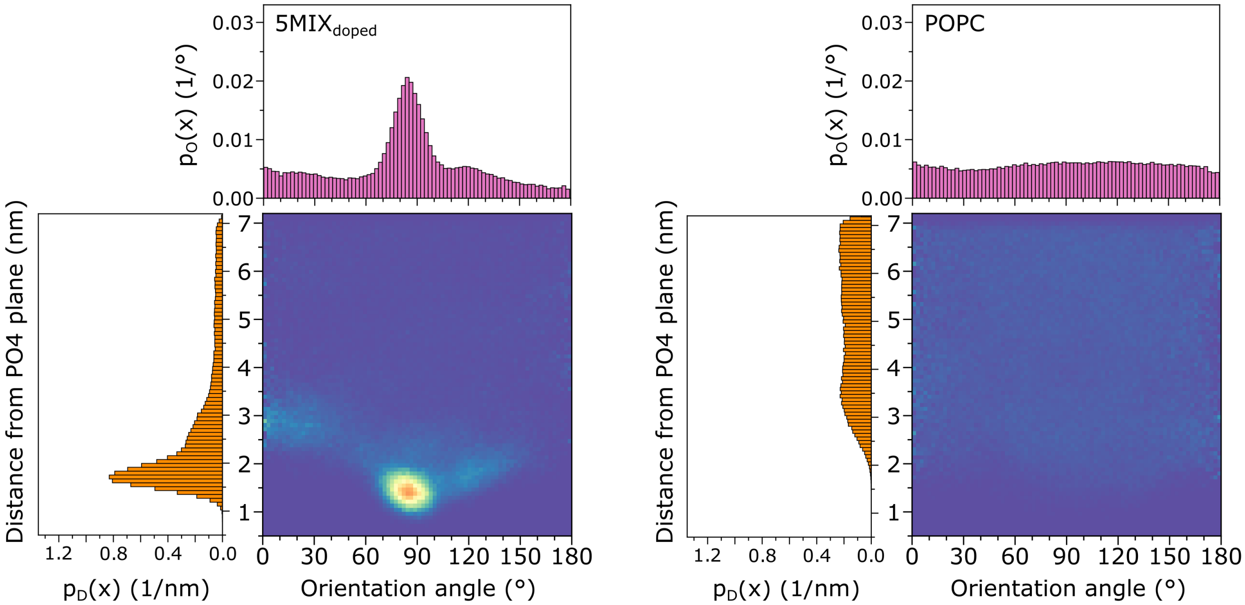


Supplementary Figure S2: **Orientation-distance population analysis of the C2C domain on the membrane surface.** The orientation-distance population profile for 5MIX$_{\text{doped}}$ lipid composition is depicted on the left, and the one for pure POPC on the right. Each 2D population is formed by combining the normalized distance and orientation angle probability densities, $p_D(x)$ and $p_O(x)$, which are presented on the left side and top of the 2D distributions, respectively. The color scale bar is identical to the one in Figure 2 of the main manuscript.

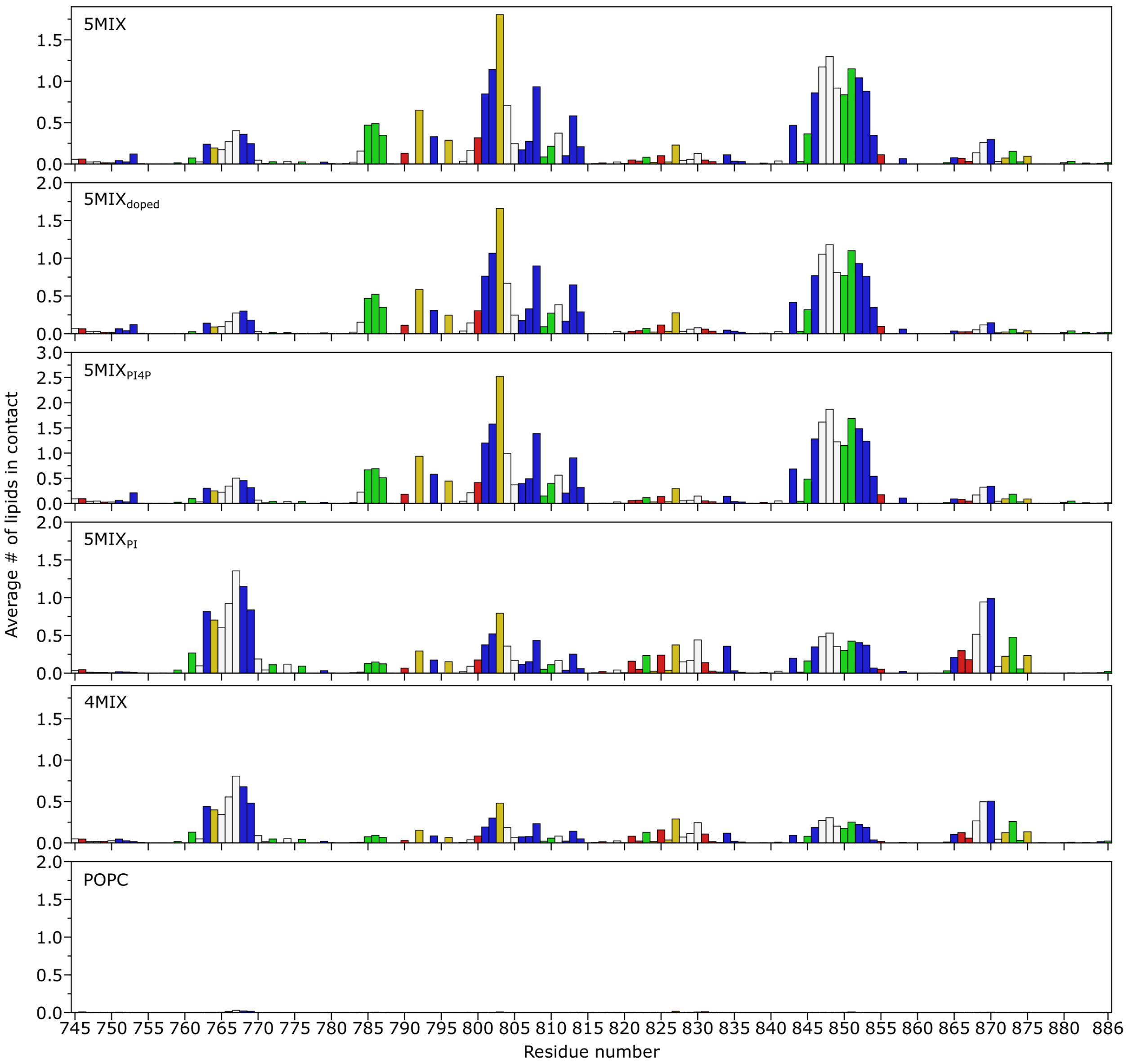


Supplementary Figure S3: **Average contacts with the bilayer per residue plotted as a function of the C2C-domain sequence for all lipid compositions.** Amino acids are colored by their chemical properties, with red representing acidic (negatively charged at physiological pH), blue for basic (positively charged at physiological pH), green for polar, white for apolar, and mustard yellow for aromatic. The first panel shows average contacts for all lipids in 5MIX, followed by 5MIX$_{doped}$, 5MIX$_{PI4P}$, 5MIX$_{PI}$, 4MIX, and POPC. Please note that the y-axis scaling is not uniform across the panels.

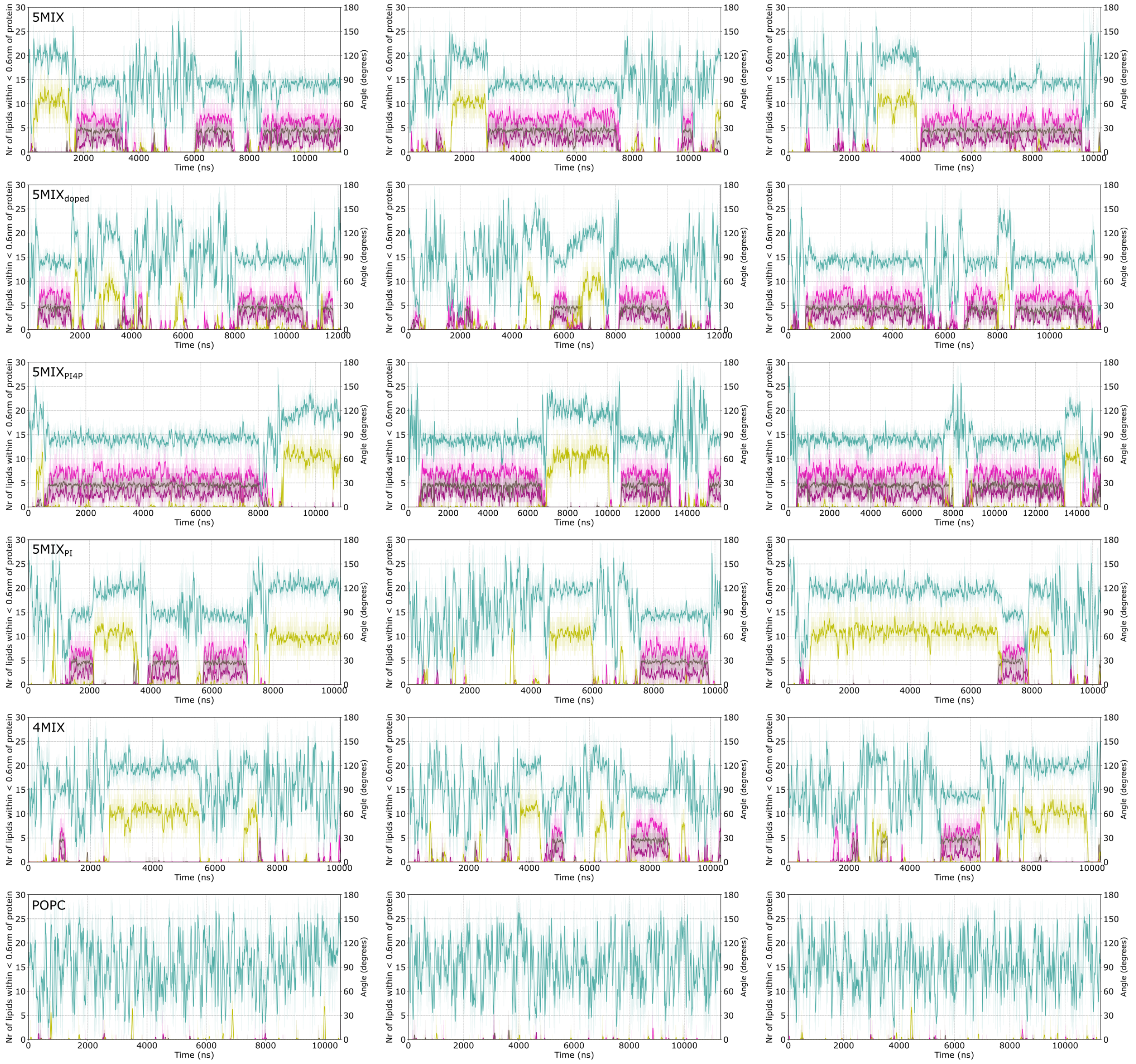


Supplementary Figure S4: **Temporal evolution of the lipid interactions of the binding sites defined in Table 1 of the main manuscript and protein orientation angle.** Each row shows the three replicas for one lipid composition: 5MIX, 5MIX$_{\text{doped}}$, 5MIX$_{\text{PI4P}}$, 5MIX$_{\text{PI}}$, 4MIX, and POPC. The transparent lines represent the raw data with 250 ps interval, while the bold lines represent the running average of the data with a 25 ns sampling window.

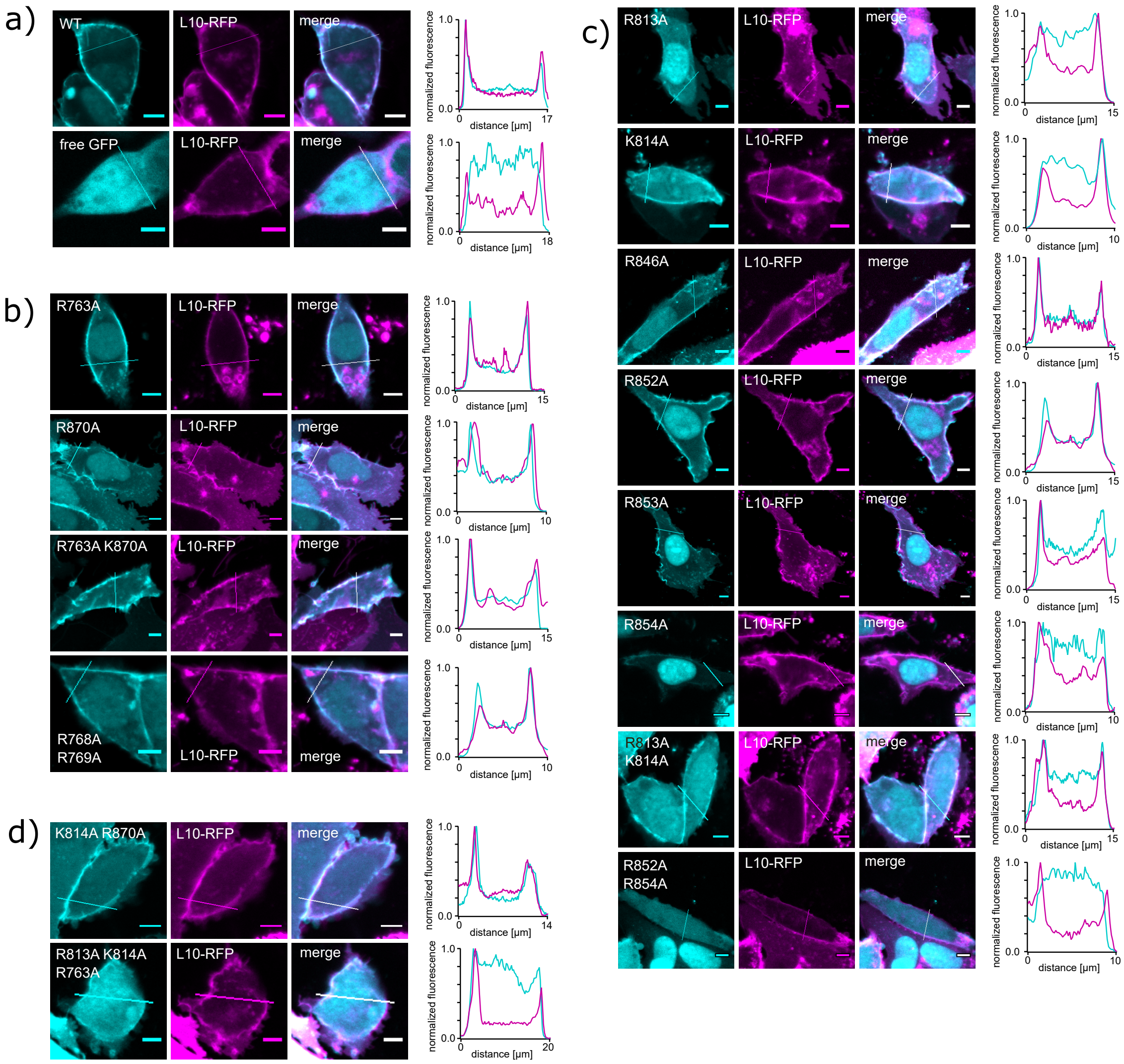


Supplementary Figure S5: **Plasma membrane localization analysis of various mutants of the C2C domain.** Representative confocal images of CHO cells co-expressing GFP-C2C domains (cyan) and the RFP-tagged PM marker L10 (magenta) are shown. Scale bars = 5 µm. Line profiles across an entire cell are highlighted in the images and shown on the right. GFP and RFP fluorescences were normalized to minimal and maximal values. a) PM localization of controls: wild-type C2C domain (WT) and free GFP. b) PM localization of mutants of the PS binding site of the C2C domain. c) PM localization of mutants of the $PIP_x$ binding site of C2C. d) PM localization of mutants of both binding sites of C2C.
